# A conformational switch in clathrin light chain regulates lattice structure and endocytosis at the plasma membrane of mammalian cells

**DOI:** 10.1101/2022.03.21.485138

**Authors:** Kazuki Obashi, Kem A. Sochacki, Marie-Paule Strub, Justin W. Taraska

## Abstract

The conformations of endocytic proteins and their interactions are key regulators of clathrin-mediated endocytosis. Three clathrin light chains (CLC), along with three clathrin heavy chains, assemble to form single clathrin triskelia that link into a geometric lattice that curves to drive endocytosis. Conformational changes in CLC have been shown to regulate triskelia assembly in solution, yet the nature of these structural changes, and their effects on lattice growth, curvature, and endocytosis in cells are unclear. Here, we develop a correlative fluorescence resonance energy transfer (FRET) and platinum replica electron microscopy method, named FRET-CLEM. With FRET-CLEM, we measure conformational changes in proteins at thousands of individual morphologically distinct clathrin-coated structures across cell membranes. We find that the N-terminus of CLC moves away from the plasma membrane and triskelia vertex as lattices curve. Preventing this conformational switch with acute chemical tools inside cells increased clathrin structure sizes and inhibited endocytosis. Therefore, a specific conformational switch in CLC regulates lattice curvature and endocytosis in mammalian cells.

## Introduction

Clathrin-mediated endocytosis is the primary internalization pathway in eukaryotic cells, and is key to many processes, including nutrient uptake, excitability, signaling, and the recycling of membrane components^1^. Over 50 proteins have been implicated in this process^2, 3^. Determining the nanoscale localizations^4, 5^, numbers^6, 7^, accumulation dynamics^8, 9^, interactions^1^, and conformations^10, 11^ of these proteins in cells is important for understanding how the endocytic machinery operates in health and disease.

Assembly and curvature of clathrin lattices are required steps to build cargo-loaded vesicles^12–14^. The basic unit of the clathrin lattice is the clathrin triskelion^15, 16^. This six protein complex is composed of three clathrin heavy chains and three smaller clathrin light chains (CLCs) that form a three-legged pinwheel (Fig S1A)^17^. The heavy chains provide the backbone, and light chains are thought to regulate the assembly of the lattice and effect its mechanical properties^16^. Work both *in vitro* and in cells has indicated its important role^18–22^, yet the structural, and functional actions of the light chains are not fully understood^16, 23^.

*In vitro*, clathrin triskelia assemble into spherical cages^13^. Clathrin light chain inhibits this assembly^24, 25^. Light chain binds to heavy chain through multiple interactions^16^. One specific interaction occurs through an acidic patch (EED) near the light chain’s N-terminus and the heavy chain’s knee (Fig S1A)^26^. This interaction prevents cage assembly by regulating the heavy chain knee conformation (Fig S1B)^17^. If CLC binds to the heavy chain through these N-terminal contacts, CLC adopts an extended conformation, stabilizing a straight conformation of the heavy chain knee. The straight conformation cannot assume the angles needed for cage assembly. However, if this interaction is prevented, the heavy chain knee is free to move, allowing the knee to bend and assemble as a cage. Thus, conformational switching of CLC is proposed to control the assembly of lattices^17^. Yet, the nature of these structural changes, and their effects on lattice growth, and curvature at the membrane of living cells are mostly unknown.

Recent super-resolution imaging studies have mapped changes in protein locations at distinct stages of endocytosis at the scale of tens of nanometers^4, 5^. Proteins are, however, regulated by sub-ten nanometer intra-and intermolecular conformational changes and binding events. To understand endocytosis, these conformational changes need to be determined^3, 13^. This is a major gap in understanding endocytosis. Yet, the precision of super-resolution imaging cannot measure structural changes at these scales. Fluorescence (or Förster) resonance energy transfer (FRET), however, can fill this gap, mapping molecular interactions and conformational changes within and between proteins^27^.

Specifically, FRET occurs when a donor fluorophore and acceptor are generally separated by less than 10 nm^28^. While FRET measurements are usually made with diffraction limited imaging, super-resolved FRET methods have been reported^29^. The resolution, labeling density, and colors possible in these experiments, however, are not able to discriminate morphological differences in small organelles such as clathrin-coated structures (CCSs). For example, past FRET analysis of the protein organization of CCSs in yeast was limited to a late stage of endocytosis trapped by drug treatments^30^. Thus, existing methods cannot readily relate changes in protein conformation generated from FRET to the morphological stages of clathrin-coated pits as they grow and curve.

To overcome this gap, we turned to correlative light and electron microscopy (CLEM)^31^. CLEM can directly map fluorescence signals to single nanoscale cellular structures visualized in paired electron microscopy (EM) images^32^. Platinum replica transmission EM (PREM) of unroofed cell plasma membranes provides a uniquely high contrast, high- resolution, and wide-field view of the inner plasma membrane of cells^33^. PREM has been combined with diffraction limited^34, 35^, super-resolution^4, 36–39^, and polarized total internal reflection fluorescence (TIRF)^40^ microscopy to investigate molecular mechanisms of endocytosis, exocytosis, and the cortical cytoskeleton. Here, we develop a correlative lifetime based-FRET (FLIM-FRET) and PREM method, named FRET-CLEM. This method allows us to measure changes in protein interactions and conformations at distances less than 10 nm at single structurally-defined organelles in single cells. We investigate the conformational changes in clathrin light chain by mapping distance changes both parallel and perpendicular to the plasma membrane. We find that the N-terminus of CLC moves away from both the CLC C-terminus and the plane of the plasma membrane as clathrin sites gain curvature. To determine the mechanistic impact of these structural changes, we develop a method to directly manipulate the N-terminal position of clathrin light chain using a chemically-inducible dimerization system. These manipulations were confirmed with FRET. With acute chemical perturbation, we show that inhibiting conformational changes in CLC’s N-terminus increases clathrin lattice size, impairs maturation of clathrin structures at the plasma membrane, and inhibits transferrin endocytosis. Together, these data reveal a new conformational switch in clathrin light chain that regulates endocytosis in living cells.

## Results

### Correlative FLIM-FRET and PREM method (FRET-CLEM)

To determine sub-ten nanometer conformational movements during endocytosis, we established a new correlative FLIM-FRET and PREM method, which we named FRET- CLEM (Fig 1). We chose monomeric EGFP and a dark yellow fluorescent protein, ShadowY^41^, as an optimized FRET pair by comparing six potential fluorescent protein (FP) pairs (Fig S2A). Calculations from the emission and absorbance spectra of this pair resulted in an estimated *R_0_* value (distance of 50% FRET efficiency) of 60 Å (Fig S2B). In line with these calculations, tandem EGFP-ShadowY probes fused to CLC showed >50% FRET efficiency on the plasma membrane (Fig S3, A-D). We used this construct as a positive FRET control. Next, to test whether FRET could be localized to single CCS, we performed FRET-CLEM measurements on HeLa cells expressing EGFP-CLC or EGFP- ShadowY-CLC (Fig 1A). First, cells were unroofed to expose the inner surface of the plasma membrane and rapidly fixed^33^. These membranes provide uniquely high contrast, low background, ultra-thin samples for fluorescence imaging^31^. Unroofed cells were then imaged with FLIM and subsequently prepared for platinum replica electron microscopy (PREM). Past work has shown that these sample preparation steps do not measurably change the morphology and structure of the membrane and its associated endocytic organelles^4, 36, 42^. This correlative method allows us to assign single diffraction-limited fluorescent spots from a FLIM image to a single clathrin structure visualized in PREM (Fig 1A). Because PREM has a high spatial resolution, single fluorescent spots in the FLIM image can be further classified according to the nanoscale structural features of the clathrin lattice (Fig 1B)^42^. We then tested whether fluorescence lifetimes can be analyzed at single CCS resolution. With the total photon number detected by our FLIM acquisition parameters (Fig S3, E-G, and Materials and Methods), fluorescence lifetime decay curves determined from single CCSs were clearly separated between negative EGFP-CLC and positive EGFP-ShadowY-CLC FRET controls (Fig 1C). Specifically, they could be fit with a bi-exponential where the FRET efficiency was determined for each CCS (Fig S3H)^43^.

**Figure 1.**
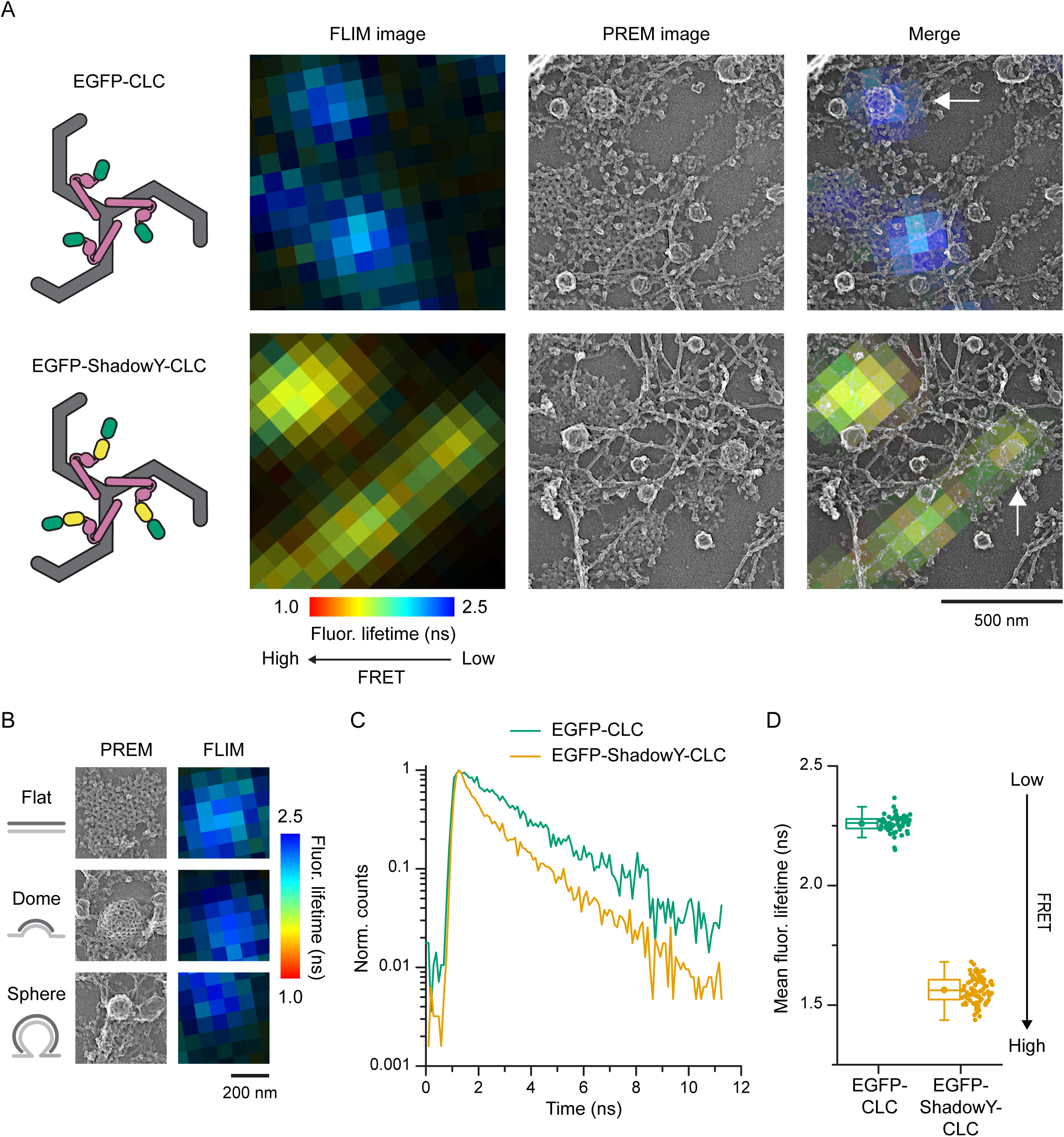
FRET-CLEM. (A) Correlative FLIM-FRET and PREM images of unroofed membranes of HeLa cells expressing EGFP-CLC (top) or EGFP-ShadowY-CLC (bottom). FLIM images (left; photon counts are represented by brightness and fluorescence lifetimes are represented by pseudo color), PREM images (center), and merge images (right). Scale 500 nm. (B) PREM images of CCSs on an unroofed membrane of a HeLa cell expressing EGFP- CLC that were classified as flat, domed, or sphere and corresponding areas from a FLIM image. Scale 200 nm. (C) Fluorescence lifetime decays from single CCSs indicated by arrows in panel A. (D) Mean fluorescence lifetimes from single CCSs on an unroofed membrane of HeLa cells expressing EGFP-CLC or EGFP-ShadowY-CLC. *n* = 58 CCSs from 1 cell (EGFP-CLC) and 81 CCSs from 1 cell (EGFP-ShadowY-CLC). For box plots, box is interquartile range, center line is median, center circle is mean, outliers are a coefficient value of 1.5.

However, curve-fitting was suboptimal for small CCSs due to the limited number of photons^44^. Thus, to measure as many clathrin structures as possible across a cell, the mean fluorescence lifetime was used to estimate FRET efficiencies across all structures (Fig 1D). These results show that FRET-CLEM can be used to generate FRET-based atomic-scale distances at single CCS resolution at morphologically distinct stages of endocytosis at the plasma membrane of mammalian cells.

### Conformational changes in CLC in cells

Next, we applied FRET-CLEM to study conformational changes in CLC at the plasma membrane. *In vitro*, clathrin triskelia assemble into empty cages and CLC regulates this assembly^26^. CLC has been proposed to take two different conformations when bound to triskelia, an extended and bent conformation (Fig 2A)^17^. Here, we assumed three models of CLC conformations in cells based on *in vitro* models (Fig 2A). First, we assume the spherical triskelia in cells resembles *in vitro* assembled cages and contains bent CLC. Next, we propose three possible models for light chain dynamics during curvature. In the first, CLC does not change conformations after assembly on the plasma membrane. Here, CLC assumes the bent conformation regardless of the curvature stage of the lattice (flat, domed, spherical). In the second, the proportions of extended and bent CLC gradually shifts as curvature increases during endocytosis. In the third, as yet undescribed conformations of CLC are present in flat and domed clathrin, that switches to the bent conformation in spheres.

**Figure 2.**
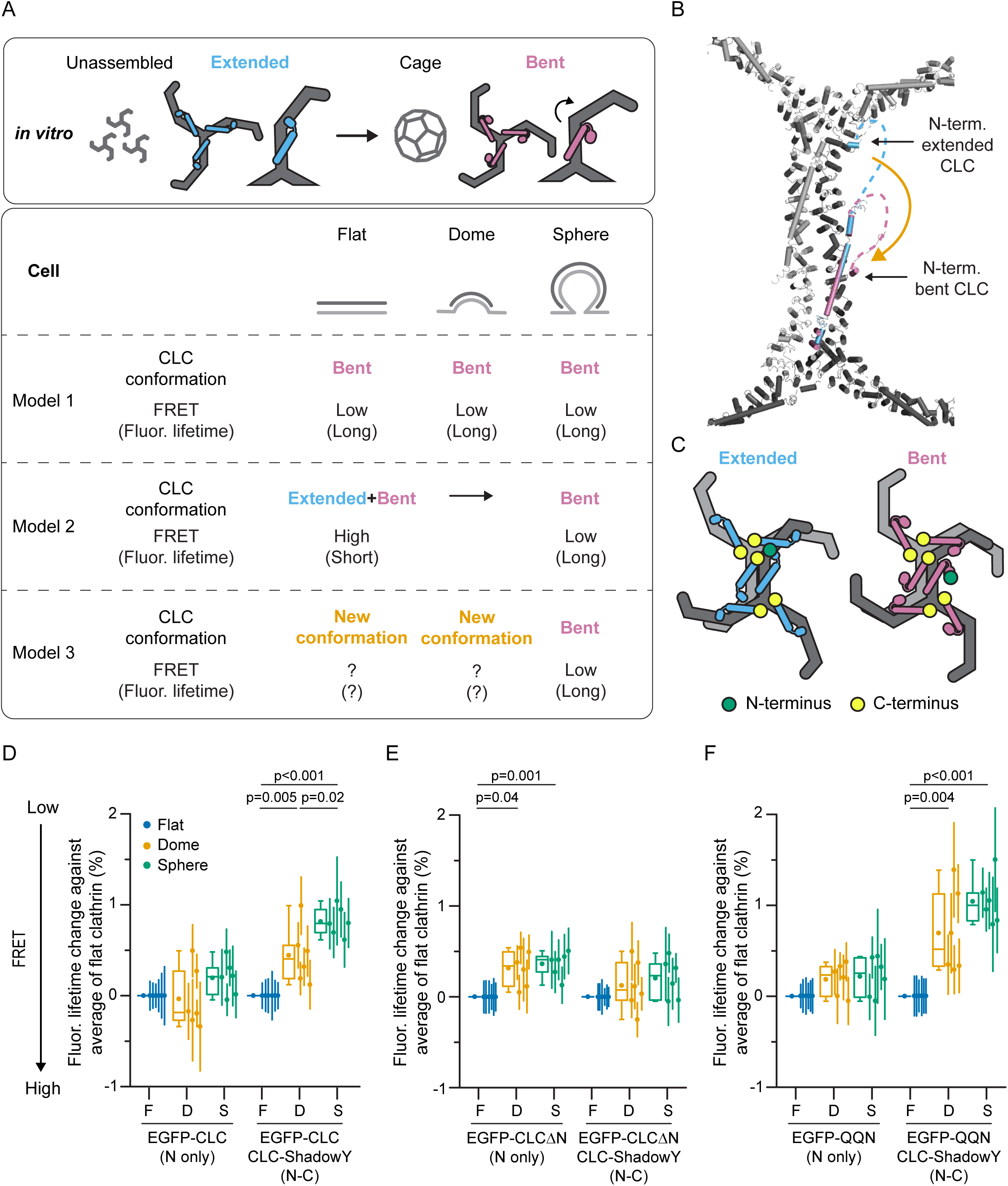
CLC conformational changes in cells. (A) Schematic models of CLC conformation at different assembly states *in vitro* or in living cells and their expected FRET efficiencies between the N- and C-terminus of CLCs. (B) A structural model with the assumption that either extended (light blue) or bent conformations (magenta) of EGFP-CLCs assemble into the lattice. The model is based on PDB 3LVG and 6WCJ. 3LVG is overlaid with 6WCJ. (C) Schematic models of two assembled triskelia with extended (left) or bent CLCs (right). A CLC N-terminus (green) and C-terminus of surrounding CLCs (yellow) are shown. (D) FRET-CLEM was performed on HeLa cells expressing either EGFP-CLC, or EGFP- CLC and CLC-ShadowY. Mean fluorescence lifetimes from single CCSs were analyzed by categorizing them according to lattice structures (flat, domed and sphere) and they were compared to the average values of flat structures. *n* = 6 cells for each condition. (E) FRET-CLEM on HeLa cells expressing either EGFP-CLCΔN, or EGFP-CLCΔN and CLC-ShadowY. *n* = 6 cells for each condition. (F) FRET-CLEM on HeLa cells expressing either EGFP-QQN (QQN mutant of CLC), or EGFP-QQN and CLC-ShadowY. *n* = 6 cells for each condition. One-way ANOVA, then Tukey’s test. Each dot is from one cell experiment and errors are SE. For box plots, box is interquartile range, center line is median, center circle is mean, outliers are a coefficient value of 1.5.

To test these models, we used FRET-CLEM. Specifically, the distances between the N-terminus and C-terminus of surrounding CLCs differ substantially between the extended and bent conformations according to structural models^17, 45^ (Fig 2, B and C, and Fig S1, C and D). Thus, if the population of extended and bent conformations change on the plasma membrane during lattice assembly, FRET between the N- and C-terminus of CLCs should likewise change (model 2 in Fig 2A). Alternatively, if the conformations do not change, FRET should remain the same across all clathrin subtypes (model 1 in Fig 2A).

To compare models 1 and 2, we performed FRET-CLEM measurements on HeLa cells expressing EGFP-CLC, or EGFP-CLC and CLC-ShadowY (Fig 2D and Fig S4, A and B). Mean fluorescence lifetimes from single CCSs were analyzed by grouping them according to lattice curvature determined from PREM images (flat, domed, and sphere; Fig 1B). In cells expressing both EGFP-CLC and CLC-ShadowY, fluorescence lifetimes increased as clathrin lattices curved (Fig 2D). This substantial increase did not occur when expressing only EGFP-CLC. The neuronal isoform of CLC which has an insertion of residues near the C-terminus^16^ and was used in the previous *in vitro* study^17^ showed similar lifetime changes (Fig S5). In living cells, FRET efficiencies between EGFP-CLC and CLC-ShadowY were higher on CCSs than in the cytosol (Fig S6). However, unlike previous measurements in solution^17^, our FRET measurements reflected both intra- and inter-triskelia FRET. Next, we measured EGFP-CLCΔN (Fig 2E and Fig S4, C and D) which is a truncation mutant of the flexible N-terminal domain (residues 1-89). In this mutant, EGFP would be bound near the heavy chain binding helix^15^. For this truncation, fluorescence lifetimes did not substantially change across different lattice states. These data indicate that the N-terminal position of CLC undergoes a conformational switch at clathrin lattices on the plasma membrane during endocytosis. These data are not consistent with model 1.

Further, fluorescence lifetimes from cells expressing EGFP-CLC and ShadowY-CLC changed across different stages (Fig S4, G and H). This supports the movement of CLC N-terminal region. Next, to further test model 2, we tested the QQN mutants of clathrin light chain (residues 20–22 were substituted from EED to QQN) (Fig 2F and Fig S4, E and F).

QQN mutants are deficient in binding to clathrin heavy chain near the triskelion vertex due to a loss of a negatively-charged patch at the N-terminus (Fig S1A)^26^. Thus, they are inhibited from adopting the proposed extended conformation^17^. Here, if model 2 is correct, FRET across the structural states should differ between wild type and QQN mutants.

Specifically, QQN mutants are expected to show a smaller displacement in conformations and smaller changes in fluorescence lifetime as the lattice curves. However, in these mutants, we found similar fluorescence lifetime changes as in the wild type protein (Fig 2 D and f). These data do not support model 2. Thus, we conclude that CLC structural movements at the plasma membrane cannot be described by current *in vitro* models (model 1 or 2).

### CLC N-terminal region moves away from the plasma membrane

Next, we measured FRET between EGFP positioned in different domains of clathrin light chain and the membrane resident dark quencher dipicrylamine (DPA) (Fig S2C)^46^. Because the plane of the membrane is fixed, FRET between EGFP and DPA provides relative distances perpendicular to the plane of the plasma membrane^47, 48^. Specifically, when EGFP sits near the membrane, FRET efficiency is high and fluorescence lifetimes are short (Fig 3A). With DPA, CLC-EGFP showed the shortest lifetime (Fig 3, B and C). This indicates that the C-terminus of CLC is located close to the plasma membrane. In contrast, EGFP-CLC showed the longest fluorescence lifetime. EGFP-CLCΔN showed an intermediate lifetime. From these data, we can position segments of CLC relative to the plane of the membrane (Fig 3D). Unlike classic models^17^, we find that the N-terminus of CLC bound to clathrin-coated structures at the plasma membrane is located farther into the cytoplasm than the proximal leg of clathrin heavy chain.

**Figure 3.**
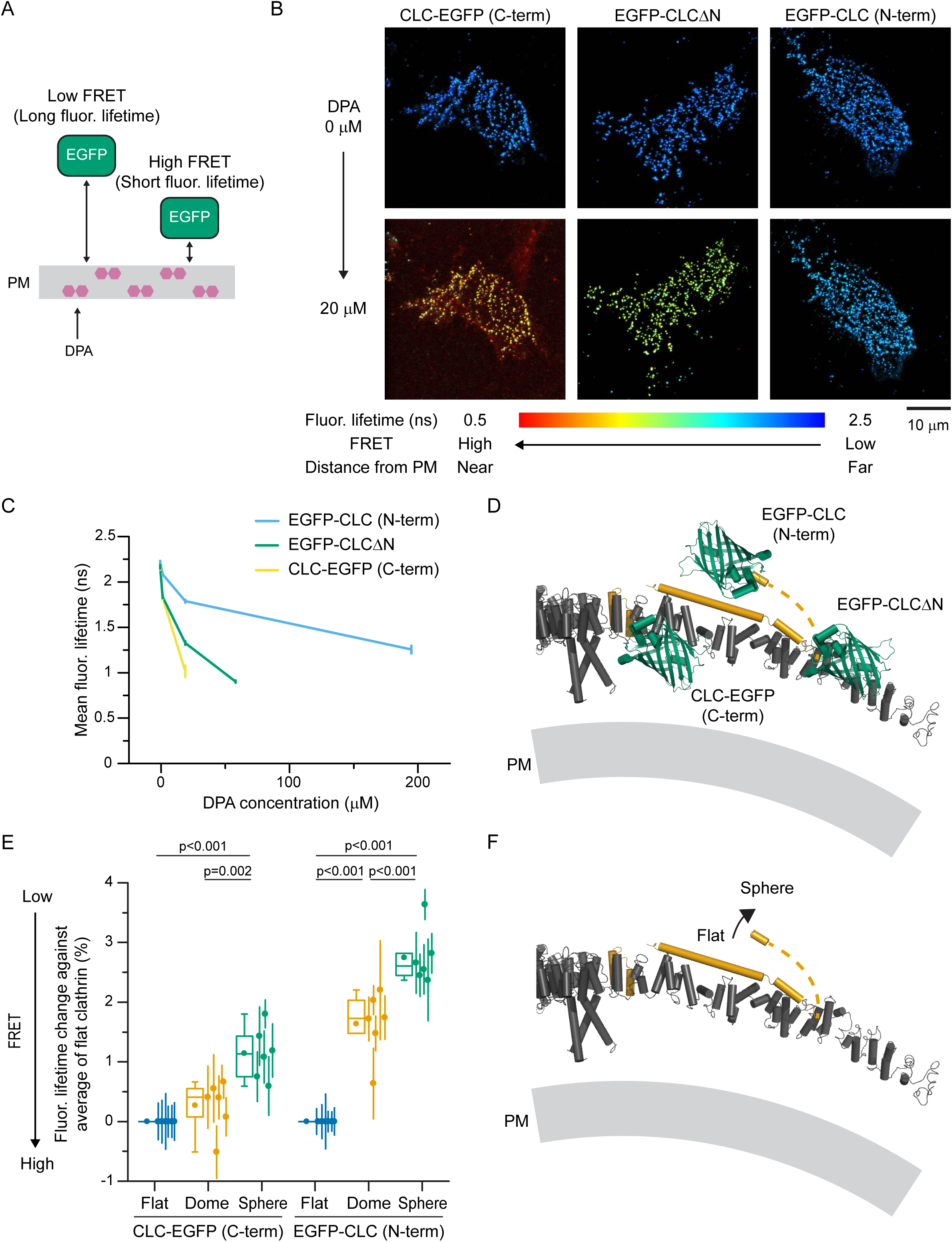
CLC N-terminal position perpendicular to the plane of the plasma membrane. (A) Dipicrylamine (DPA) is a nonfluorescent hydrophobic anion that incorporates into membranes. DPA quenches EGFP in a distance dependent manner by FRET. (B) FLIM images of an unroofed membranes of HeLa cells expressing CLC-EGFP (left), EGFP-CLCΔN (center), or EGFP-CLC (right) without (top) or with 20 μM DPA (bottom). Scale 10 μm. (C) Mean fluorescence lifetimes with different DPA concentrations. *n* = 10 (EGFP- CLC), 11 (EGFP-CLCΔN), and 8 cells (CLC-EGFP) from 2 experiments. Errors are SE. (D) A structural model of EGFP positions of CLC probes predicted from the EGFP-DPA FRET experiments. The model is based on PDB 4KW4 and 3LVG. (E) FRET-CLEM on HeLa cells expressing either CLC-EGFP or EGFP-CLC with DPA. DPA concentrations were 3 μM for CLC-EGFP and 80 μM for EGFP-CLC to obtain ∼50% FRET efficiencies to make the degree of fluorescence lifetime changes similar. Mean fluorescence lifetimes from single CCSs were analyzed by categorizing them according to lattice structures and compared to average values of flat structures. *n* = 6 cells for each condition. Each dot is from one cell experiment and errors are SE. One-way ANOVA, then Tukey’s test. For box plots, box is interquartile range, center line is median, center circle is mean, outliers are a coefficient value of 1.5. (F) A proposed structural model of CLC conformational changes predicted from both FRET-CLEM with EGFP-ShadowY and EGFP-DPA. The N-terminus of CLC moves away from both the CLC C-terminus (triskelion vertex) and the plane of the plasma membrane as clathrin lattices curve. The structural model is based on PDB 3LVG.

To investigate conformational changes during lattice curvature, we performed FRET- CLEM measurements with DPA and compared those measurement to morphological changes in clathrin lattice (Fig 3E and Fig S7). In these experiments, fluorescence lifetimes of EGFP-CLC increased as lattices curved, and the degree of change was larger than those for CLC-EGFP. This is consistent with a model where the N-terminal position of CLC changes during curvature. Thus, the combined results from EGFP-ShadowY and EGFP-DPA FRET indicate that the N-terminus of CLC extends away from the CLC C-terminus (triskelion vertex) and the plane of the plasma membrane during endocytosis (Fig 3F).

### Conformational changes in CLC regulate lattice structure and endocytosis

Next, we tested whether these conformational changes in CLC regulate clathrin-mediated endocytosis. To manipulate the CLC N-terminal position, we used a rapamycin inducible FKBP/FRB dimerization system^49^. Specifically, we employed the T2098L mutant of FRB which is heterodimerized to FKBP by the rapamycin analog, AP21967^50^. We attached FKBP to the N-terminus of CLC, and one or two FRBs (FRB×2) to the C-terminus of either the membrane-bound PH domain from PLCδ1 (PH)^51^ (Fig 4A) or epsin1^52^ (Fig S8A). We made both FRB and FRB×2 probes for the PH domain to extend the reach of the system.

**Figure 4.**
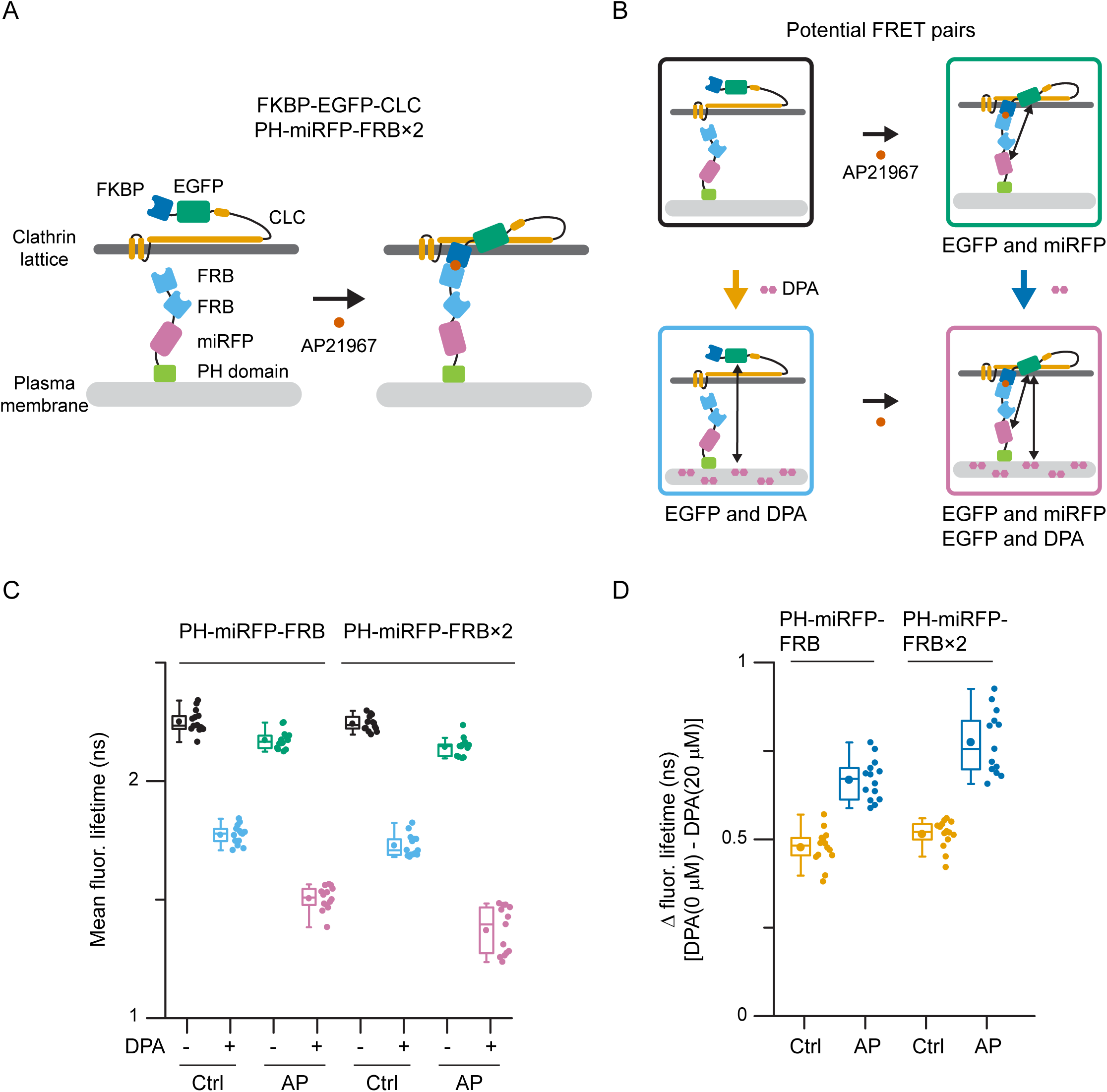
Manipulation of CLC N-terminal position using a chemically inducible dimerization system. (A) Schematic models of the chemically inducible FKBP/FRB dimerization system. FKBP is attached to the N-terminus of CLC, and two FRBs (FRB×2) are attached to the C-terminus of PH domain from PLCδ1. A rapamycin analog, AP21967, induces heterodimerization between FKBP and the T2098L mutant of FRB. (B) Potential FRET pairs without or with AP21967 treatment either absence or presence with DPA. (C) Fluorescence lifetime measurements were performed on unroofed membranes of HeLa cells expressing FKBP-EGFP-CLC either with PH-miRFP-FRB, or PH- miRFP-FRB×2 without or with 20 μM DPA. Cells were unroofed after 15 min incubation with AP21967 or ethanol (control). *n* = 14 (PH-miRFP-FRB, control), 14 (PH-miRFP-FRB, AP21967), 14 (PH-miRFP-FRB×2, control), and 13 cells (PH- miRFP-FRB×2, AP21967) from 2 experiments. (D) Differences in fluorescence lifetimes without and with DPA.

To confirm that the N-terminal position of FKBP-EGFP-CLC changes relative to the plasma membrane after dimerization, we measured EGFP fluorescence lifetimes without or with AP21967 in the presence of DPA (Fig 4, B and C, and Fig S8B). Changes in the position of EGFP can be estimated by comparing the difference in fluorescence lifetimes between the control and DPA treated membranes (Fig 4D and Fig S8C). For all three probes, differences in EGFP lifetimes with AP21967 treatment were larger than those without AP21967. These results indicate that the CLC N-terminus moves towards the plasma membrane as a result of FKBP/FRB dimerization. Because PH-miRFP-FRB×2 showed larger lifetime change (Fig 4D) and stronger accumulation with AP21967 than PH-miRFP-FRB (Fig S9A), we used PH-miRFP-FRB×2 and epsin1-miRFP-FRB (or PH-mCherry-FRB×2 and epsin1-mCherry-FRB) to alter the structure of clathrin light chain in living cells.

First, using our FKBP/FRB constructs, we investigated the impact of moving the CLC N-terminus towards the plasma membrane on the structure of clathrin lattices^42^. Here, unroofed membranes from cells expressing FKBP-EGFP-CLC and either PH-miRFP- FRB×2 or epsin1-miRFP-FRB and treated without or with AP21967 were imaged with PREM (Fig S10A). Unlike the previous homodimerization system^53^, clathrin lattices were not noticeably distorted. The average size of all visible flat and domed clathrin structures increased with AP21967 treatment for PH-miRFP-FRB×2 expressing cells [31463 ± 859 nm^2^ (ctrl) and 42919 ± 2278 nm^2^ (AP) for flat, 29821 ± 576 nm^2^ (ctrl) and 36471 ± 2862 nm^2^ (AP) for domed, 15311 ± 588 nm^2^ (ctrl) and 17303 ± 691 nm^2^ (AP) for sphere] (Fig 5, A-C).

**Figure 5.**
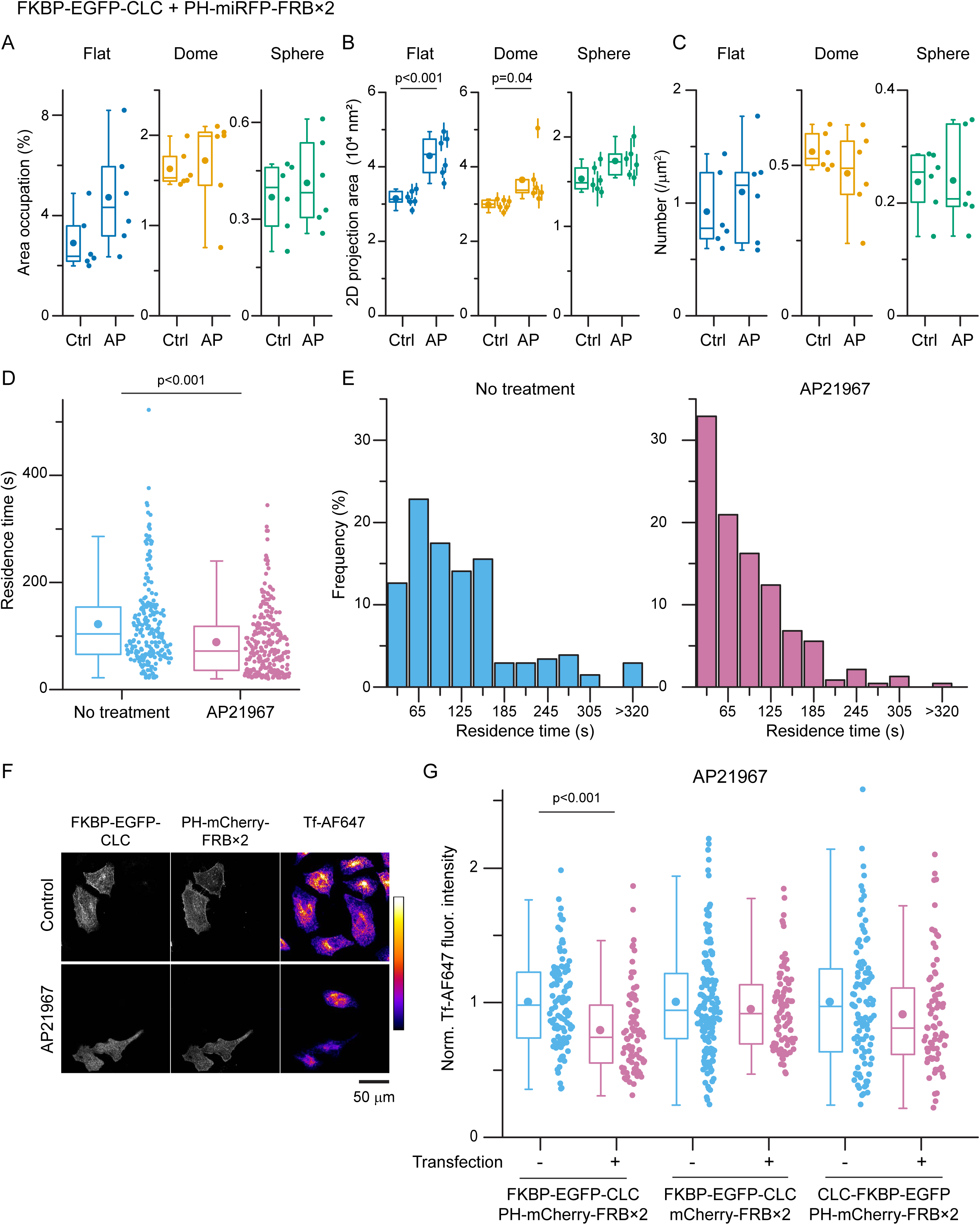
Manipulation of CLC conformation changed lattice structures, dynamics, and endocytosis. (A-C) Unroofed membranes from cells expressing FKBP-EGFP-CLC and PH-miRFP- FRB×2 treated with AP21967 or ethanol (control) were imaged with PREM. Two- dimension area of single CCS were manually segmened and measured. Membrane area occupation against the total analyzed membrane area (A), two-dimension projection area (B), and density (C) of flat, domed, and sphere CCSs were compared. Each dot is from one cell experiment, and errors are SE. *n* = 6 cells for each condition. The average measured area / cell (mean ± SE) = 213 ± 34 (control) and 165 ± 10 μm^2^ (AP21967). (D) Live cell time lapse TIRF imaging on HeLa cells expressing FKBP-EGFP-CLC and PH- miRFP-FRB×2 without or with AP21967 treatment. Tracks with over 20 s were analyzed and residence times were compared. *n* = 206 spots from 5 cells (no treatment) and 234 spots from 5 cells (AP21967). An unpaired t test was used. (E) Histogram of residence times. (F) Confocal projection images of Alexa Fluor 647 conjugated transferrin (Tf-AF647) uptake in HeLa cells expressing FKBP-EGFP-CLC and PH-mCherry-FRB×2 treated with AP21967 or ethanol (control). Fluorescence intensity of Tf-AF647 is represented by pseudo color. Scale 50 μm. (G) Transferrin uptake in HeLa cells expressing FKBP and FRB probes treated with AP21967. Fluorescence intensities of incorporated Tf-AF647 normalized by non- transfected cells in the same sample were compared between non-transfected and transfected cells. *n* = 75-142 cells from 2 experiments for each conditions. For box plots, box is interquartile range, center line is median, center circle is mean, outliers are a coefficient value of 1.5. An unpaired t test was used.

Although expression of the epsin1-miRFP-FRB decreased the average clathrin-coated structure size on its own, AP21967 treatment increased flat and domed clathrin areas similar to the extent seen with the PH domain-based tethers (Fig S10, B-D). We conclude that manipulation of the CLC N-terminal position directly effects the structure of clathrin.

Next, to investigate how changing CLC conformations effects CCS assembly and maturation, we performed live cell time lapse evanescent field imaging. Cells expressing FKBP-EGFP-CLC and either PH-miRFP-FRB×2 (Fig 5, D and E) or epsin1-miRFP-FRB (Fig S11) were imaged with TIRF in the absence or presence of AP21967. To quantitate CCS dynamics, we analyzed the residence times of FKBP-EGFP-CLC spots on the plasma membrane. For PH-miRFP-FRB×2, the residence times became shorter after AP21967 addition (Fig 5, D and E, and Fig S11B). Similarly, although expression of epsin1-miRFP- FRB shorten the residence times slightly, AP21967 treatment further shorten them (Fig S11). These data indicate that tethering the CLC N-terminal position towards the plasma membrane likely facilitates the disassembly of CCSs.

Finally, we investigated whether manipulation of the CLC N-terminal position effects transferrin endocytosis in whole cells (Fig 5, F and G and Fig S12). As a control for binding of FRB to the CLC N-terminus, a cytosolic probe, mCherry-FRB×2, was used.

Furthermore, as a control of clathrin tethering to the plasma membrane, and also FRB accumulation on CCSs, a C-terminal-linked FKBP probe, CLC-FKBP-EGFP, was tested. For all five probe pairs, FKBP/FRB dimerization was confirmed by clustering of FRB at CCSs (Fig S9B) and/or changes in FRET efficiency between EGFP and mCherry (Fig S9C). Also, expression of these probes did not change transferrin uptake without AP21967 treatment (Fig S12B). We found that transferrin uptake decreased with AP21967 for PH– mCherry-FRB×2 but not for the control mCherry-FRB×2 (Fig 5G). Further, for C-terminus-attached FKBP control (CLC-FKBP-EGFP), PH-mCherry-FRB×2 did not change transferrin uptake (Fig 5G). Although transferrin uptake of epsin1-mCherry-FRB expressing cells was decreased for both CLC N- and C-terminus-attached FKBP probes, the degree of change with the C-terminus-attached FKBP was smaller than that for the N-terminus-attached FKBP (Fig S12C). These results indicate that although epsin1 had some effect on endocytosis, likely driven by epsin’s roles in endocytosis, manipulation of the CLC N-terminal position inhibited transferrin uptake. These data are consistent with the decrease in the residence time of CCSs we observed in live cell imaging. From these collective data, we concluded that the conformational changes in CLC we mapped in FRET-CLEM experiments are key structural changes required for endocytosis of cargo-loaded vesicles in mammalian cells. Thus, the movement of the N-terminal domain away from the clathrin vertex and plasma membrane is a key regulatory step in clathrin-mediated endocytosis.

## Discussion

We have explored the molecular-scale conformational changes in clathrin light chain at single clathrin sites at the plasma membrane of mammalian cells. We find that the N-terminus of CLC moves away from both the triskelion vertex and the plasma membranes as clathrin lattices curve. Blocking this movement increased clathrin lattice sizes at the plasma membrane and reduced transferrin endocytosis. Thus, specific structural changes in CLC in clathrin lattices at the plasma membrane are regulators of curvature and endocytosis in living mammalian cells.

Conformational changes in CLC have been proposed to regulate clathrin assembly^17^. We mapped the specific changes in cells by measuring vectors in two orthogonal axes. First, EGFP-DPA FRET measurements positioned the light chain relative to the plane of the membrane (Fig 3D), and reported the movement along the axis perpendicular to the plasma membrane (Fig 3F). Unlike classic *in vitro* models^17^, we find that the N-terminus of CLC is located farther into the cytoplasm than the proximal leg of clathrin heavy chain and moves away from the membrane. With intermolecular FRET between EGFP and ShadowY (Fig 2) at the N- and C-terminus of the light chain, we could map structures along the axis parallel to the plasma membrane. Here again, we detected a displacement of the N-terminal domain away from the CLC C-terminus (clathrin lattice vertex).

Combined, these data indicate that the N-terminus of CLC moves away from both the clathrin vertex and the plane of the plasma membrane as lattices gain curvature.

Although the exact position of the N-terminus in relation to the heavy chain proximal leg domain was not determined, our experiments could be used to provide additional insights. Specifically, we modeled how FRET efficiencies between EGFP-CLC/CLC-ShadowY (Fig S4B) or EGFP-CLC/ShadowY-CLC (Fig S4G) would change as the N-terminal position changes according to the x-ray structures^17^ across all possible spatial positions (Fig S13A). Although the expression level of ShadowY could affect the overall gross efficiency of FRET, the positional dependency, and direction of change, would not change with expression levels according to our calculations. Because FRET efficiencies were largely similar for both sites, the N-terminus of CLC is predicted to rest in the overlap regions that match these paired distances (Fig S13B, magenta circles). To accommodate these constraints, CLC would assume a slightly folded conformation (orange and green in Fig S13C) rather than a stretched conformation (blue in Fig S13C). Thus, we propose a new model of CLC conformational changes (Fig 6). Here, CLC assumes an extended and tightly bound conformation in unassembled triskelia in the cytoplasm similar to that seen in x-ray crystal structures and measured with FRET^17^. Next, when CLC assembles at the membrane as a flat lattice, the light chain changes conformations from this extended position into a new folded conformation. The N-terminus is then displaced deeper into the cytosol as clathrin lattices curve into vesicles. In support of our data, the average position of the CLC N-terminal domain is not visible by cryo-EM in purified clathrin-coated vesicles^45^. This suggests that the N-terminal domain is flexible and positioned away from the lattice in highly curved structures.

**Figure 6.**
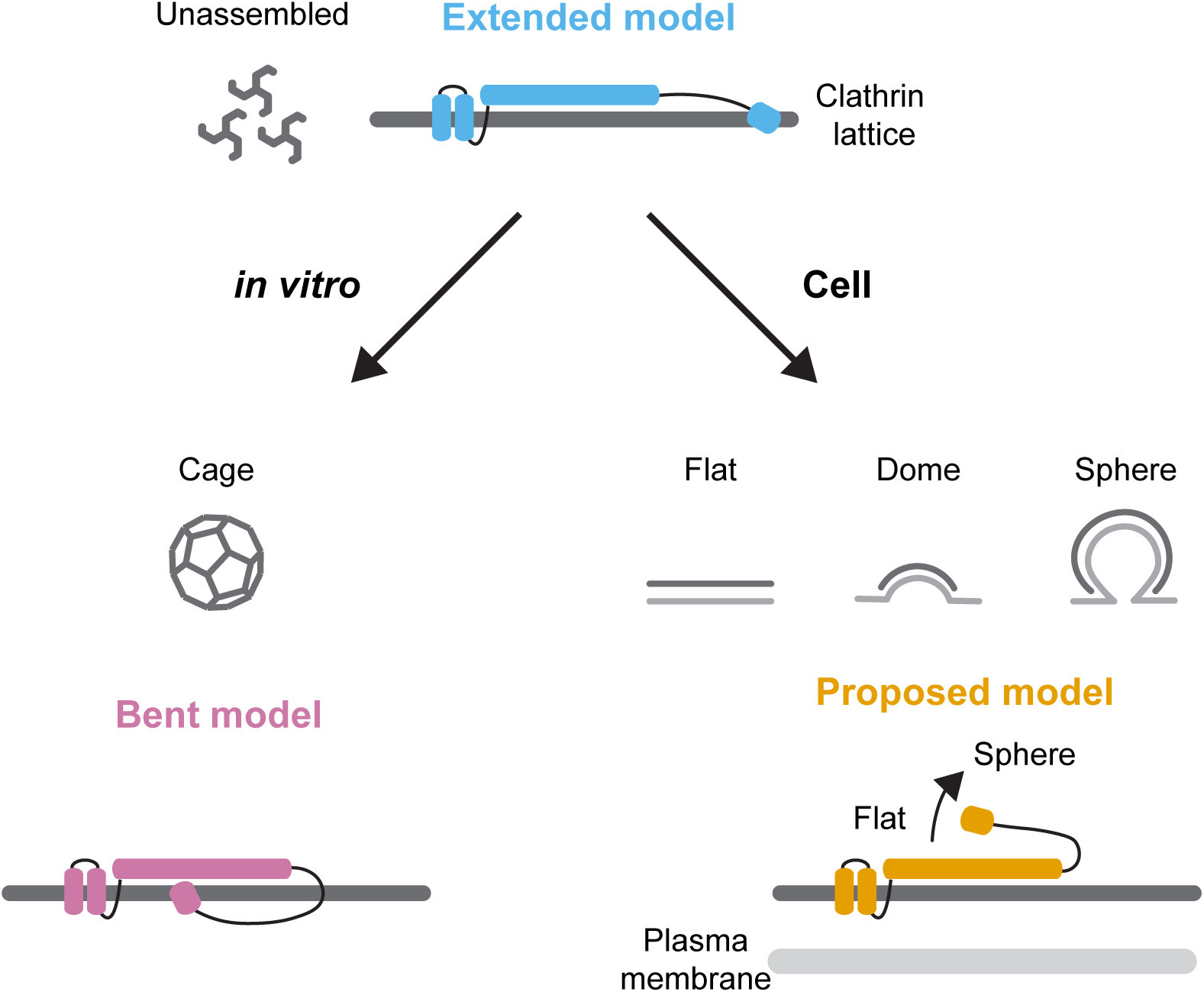
Models of conformational switch in CLC in living cells. Schematic model of conformational switch in CLC at the plasma membrane in cells. CLC assumes an extended conformation in unassembled triskelia in the cytoplasm similar to that seen in x-ray crystal structures. Next, when CLC assembles at the membrane as a flat lattice, CLC changes conformations from the extended to a new folded conformation. The N-terminus is then displaced deeper into the cytosol as clathrin lattices curve into vesicles.

What initiates this structural change in the light chain? HIP and HIP1R interact with the CLC N-terminal residues^54, 55^. Although CLC QQN mutants are deficient in binding HIP1 and HIP1R^55^, we found that FRET between the N- and C- terminus of the CLC QQN mutant showed FRET changes comparable to wild type proteins (Fig 2, D and F). Thus, if this region is necessary for binding, HIP1R interactions are unlikely to drive these conformational changes. The roles of other proteins^16, 23^ or phosphorylation^20^ are alternate mechanisms that could drive this change. Finally, non-specific physical mechanisms such as crowding and cargo binding could induce the conformational switch^2, 13^.

*In vitro* studies have shown that CLC increases the rigidity of clathrin lattices on a solid substrate^19^. These effects were different between different CLC isoforms^22^. In living cells, CLC has been shown to be important for endocytosis under conditions of high membrane tension^18, 21^. Here, our FKBP/FRB experiments demonstrate that conformational changes in CLC regulate the structure of clathin (Fig 5 A-C), the dynamics of single clathrin sites in live cells (Fig 5 D and E), and the endocytosis of cargo in whole cells (Fig 5F and G). One possible interpretation is that the lattice can grow but not fully curve due to the inhibition of the CLC conformation switch. Possibly, clathrin lattices grow larger because treatment disturbed the balance between lattice assembly and curvature^12^. These irregularly-assembled CCSs are then disassembled by proofreading mechanisms^3^. The CLCs role in endocytosis differs between isoforms, cargos, and cell types^18, 20–22, 56–59^ and the physiological function of CLC are not fully understood^16, 23^. Clarification on how CLC conformations are regulated across isoforms and cargos will provide a unified view of the light chain’s mechanistic roles across different cells, tissues, and states.

We used a PH domain (Fig 4) and epsin1-linked FRB proteins (Fig S8) to manipulate the position of the CLC N-terminus. Both constructs changed the N-terminal position of CLC and modified the structure and dynamics of CCSs (Figs 5, S10, S11 and S12). The directions of change were similar. However, expressing epsin1-miRFP-FRB measurably decreased the size of CCSs (Figs 5B, S10C, and S14C) and shortened the residence time (Fig S11). This structural change is consistent with previous studies indicating membrane bending by molecular crowding of epsin1’s disordered domains^60, 61^. Thus, attaching miRFP-FRB to a disordered domain might facilitate membrane bending in cells. Changes in clathrin lattice size by interactions between FKBP-EGFP-CLC and epsin1-miRFP-FRB might be induced by both inhibition of CLC-conformational changes and structural changes in disordered domains of epsin1. Future work is needed to disentangle these effects.

Our work has some specific limitations. For example, we transiently expressed FP-tagged probes to measure FRET. Thus, the expression ratio between endogenous and FP-labeled proteins could affects the overall FRET efficiency (Fig S1D and S13A). This, along with other photophysical issues, makes it difficult to convert FRET efficiencies to absolute atomic distances^62^. However, the differences in expression level do not affect the direction of FRET changes caused by conformational changes in this system (Fig S1D and S13A). In the future, dual-tagging all alleles of the endogenous protein is a direction to further refine these measurements^30^. Furthermore, the size of fluorescent proteins, while relatively small compared to the size of the clathrin complex and approaching the size of large red organic dyes, may affect a protein’s structure. Likewise, larger probes can limit the ability to detect small or distant conformational changes^62^. Thus, the incorporation of smaller tags or artificial amino acids are future directions to improve these measurements^63, 64^. Indeed, smaller probes have been shown to better recapitulate absolute distance changes seen during conformational transitions and might have less of an influence on the proteins themselves^28^. In the case of clathrin light chain, however, fusion of fluorescent proteins has not been seen to perturb endocytosis or change clathrin lattice structures^65^.

FRET efficiencies are determined by photophysical properties (excitation, emission, absorbance), distance, and orientation of the probes^27, 43^. Here, we used a flexible linker to connect the FPs to CLC. Thus, it is reasonable to propose that the population of FPs in a CCS can assume many possible orientations and the effect of an orientation bias would be small. However, we cannot exclude the small possibility that the curvature of clathrin lattices causes differences in the orientations between the donor and acceptor, which may affect the overall FRET efficiencies. However, the fact that the N-terminal changes was observed in both two FP-FRET experiments and FP-DPA FRET experiments, supports the idea that orientation effects are not large or dominant. Combined, these caveats make it difficult to directly equate FRET efficiencies to atomic distances. This is commonly the case in FRET studies done within cells^62^.

For cells expressing only EGFP probes, although there were no substantial change in fluorescence lifetimes among different lattice structures, clathrin vesicles tended to show slightly higher lifetimes than flat clathrin structures (Fig 2). Because Homo-FRET does not change the overall fluorescence lifetime^43^, this might be caused by a difference in the local environment^66^, or a systematic counting error due to the differences in the photon count rate among CCSs with different fluorophore densities^67^. These effects, however, were small and consistent across samples.

We developed a new correlative FLIM-FRET and PREM method to track the conformational changes in CLC at single sites of endocytosis in cells. The N-terminus of CLC makes a dramatic movement away from the clathrin lattice vertex and deeper into the cytosol as lattices curve. These conformational dynamics are key for clathrin-mediated endocytosis. These data, combined with the rich biochemical, functional, genetic, and biophysical information on membrane traffic, will lead to a more robust understanding of how endocytic proteins work together during clathrin-mediated endocytosis to drive the internalization of cargo. More generally, our new method maps molecular interactions and conformational changes at targeted proteins at identified sites in the complex environment of the cell’s plasma membrane. Thus, FRET-CLEM can be used to investigate conformational changes of any accessible membrane-associated protein including ion channels, transporters, receptors, adhesion proteins, or enzymes, in the context of their local plasma membrane environments^28, 31^.

## Materials and Methods

### Cell culture

HeLa cells (ATCC, CCL-2) were maintained at 37°C, with 5% CO_2_ in DMEM (Gibco, 11995073) supplemented with 10% fetal bovine serum (ATLANTA biological, S12450H) and 1% vol/vol penicillin/streptomycin (Invitrogen, 15070-063). The cells were placed on poly-L-lysine coated coverslips (Neuvitro, GG-25-1.5-PLL). They were transfected with 0.75 mL of Opti-MEM (Life Technologies, 31985062), 3.8 μL Lipofectamine 2000 (Life Technologies, 11668027), and 1.5 μg of DNA for 3 h after being introduced to the cells. Then, transfected cells were incubated in DMEM growth medium for 20-30 h before unroofing or imaging. The cells were tested negative for mycoplasma contamination.

### Plasmids

ShadowY (#104621), mScarlet-I (#85068), mTurquoise2 (#54842), miRFP703-CLCb (#79997), CFP-FKBP (#20160) were purchased from Addgene. CLCa-GFP was kindly donated by Dr. W. Almer (Oregon Health & Science University). Lyn-GFP-FRB was kindly donated by Dr. T. Inoue (Johns Hopkins University). mNeonGreen was kindly donated by Dr. J. Shah (Harvard University). mCherry-PH and epsin1-dGFP were from our previous studies^4, 68^. All plasmids used in this study were constructed using either Q5 Site Directed Mutagenesis Kit (New England Biolabs, E0554S) or In-Fusion HD Cloning Plus (Clonetech, 638920) following manufacturer’s instructions. N- and C-terminal disordered regions of FPs were deleted. And proteins of interest and either FPs, FKBP, or FRB were connected with a short flexible linker (GGSGGS). All plasmids were confirmed by sequencing (Psomagen) and identified as in Table S1.

### Fixing and unroofing

Cells were rinsed in intracellular buffer (70 mM KCl, 30 mM HEPES maintained at pH 7.4 with KOH, 5 mM MgCl_2_, 3mM EGTA), and manually unroofed with 19-gauge needle and syringe using 2% paraformaldehyde (Electron Microscopy Sciences, 15710) in the intracellular buffer. After unroofing, the coverslips were transferred to fresh 2% paraformaldehyde in the intracellular buffer for 20 min. They then washed with phosphate-buffered saline (PBS). Imaging was performed in PBS (pH 7.4) (Quality Biological, 114-058-131).

For FKBP/FRB dimerization experiments, cells were unroofed after 15 min incubation with 500 nM AP21967 (500 μM stock in ethanol) (Takara, 635055) or 0.1% w/v ethanol (control) in DMEM growth medium.

### Imaging of unroofed plasma membranes for CLEM

Unroofed cells were stained either with 16.5 pmol of Alexa Fluor 350-phalloidin (Life Technologies, A22281), Alexa Fluor 568-phalloidin (Life Technologies, A12380), or Alexa Fluor 647-phalloidin (Life Technologies, A22287) for 15 min depending on spectra of expressing FPs. Then cells were rinsed with PBS. 1 mm × 1 mm large montage was generated for proteins of interest and phalloidin using a Nikon Eclipse Ti inverted microscope with a 100×, 1.49 NA objective (Nikon, SR HP Apo TIRF) and an Andor iXon Ultra 897 EM-CCD camera under the control of Nikon Elements software. Images were obtained by TIRF illumination except for Alexa Fluor 350 which was imaged by epi illumination. This map was used to find the cells expressing the target proteins for FLIM and CLEM analysis. The imaged area was marked with a circle (4 mm in diameter) around the center of the imaged area using an objective diamond scriber (Leica, 11505059)^39, 69^. The immersion oil was carefully removed from the bottom of the glass coverslip. The sample was subsequently imaged by FLIM or stored in 2% glutaraldehyde (Electron Microscopy Sciences, 16019) at 4°C until EM sample preparation.

### FLIM

Time-domain fluorescence lifetime imaging was performed with a Leica Falcon SP8 confocal microscope with a 63×, 1.40 NA oil immersion objective (Leica, HC PL APO CS2). As first, a large image montage around the region of interest on the coverslip marked by a diamond scriber was acquired to find the cells which were identified in the TIRF montage. Then, FLIM images were collected at a lateral spatial resolution of 80 nm per pixel and with a scan speed at 3.16 μs per pixel. The confocal aperture was set at a diameter of 168 μm (2 AU @510 nm). EGFP was excited by 488 nm and 498-560 nm fluorescence was collected. A notch filter for 488 nm was used. To reduce the impact of pile-up effect, an excitation power was set to keep peak count per pulse as less than 0.1. For CLEM, the imaged sample was then stored in 2% glutaraldehyde at 4°C until EM sample preparation.

To select a FP pair for FLIM-FRET, we made tandemly connected FPs and compared their FRET efficiency (Fig S2A). Based on its high FRET efficiency and compatibility for our optical systems, we selected monomeric EGFP and ShadowY (Fig S2B). To check the relationship between the number of frames and photobleaching or FRET efficiency change, fluorescence intensity and FRET efficiency of EGFP-ShadowY-CLC was compared with various frame numbers (Fig S3, E and F). Fluorescence intensity slightly increased and FRET efficiency slightly decreased as the frame number increases. This might be due to photobleaching of ShadowY occurs faster than EGFP. Although the magnitude of ShadowY photobleaching should be smaller for lower FRET efficiency situation like intermolecular FRET, we used 150 frames to minimize the effect of photobleaching. With 150 frames, ∼10,000 photon counts per single CCS were obtained in average from a cell with medium level expression (Fig S3G). With this range of photon counts, FRET efficiency (*E*) of EGFP- ShadowY-CLC from single CCS could be estimated by fitting a fluorescence decay curve with a bi-exponential function convolved with the Gaussian pulse response function^70^ (Fig S3H).

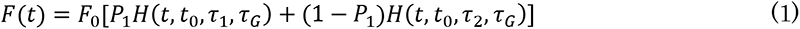

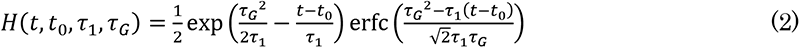

where *F*(*t*) is the fluorescence lifetime decay curve, *F_0_* is the peak fluorescence before convolution, *P_1_* is the fraction of the first component, *τ_1_* and *τ_2_* are the fluorescence lifetime of first and second components, *τ_G_* is the width of the Gaussian pulse response function, *t_0_* is time offset, and erfc is the complementary error function.

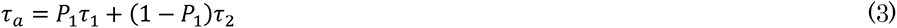

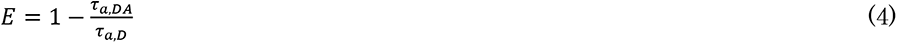

where *τ_a_* is amplitude-weighted fluorescence lifetime, *τ_a,D_* is amplitude-weighted fluorescence lifetime of the donor without acceptor, and *τ_a,DA_* is amplitude-weighted fluorescence lifetime of the donor in presence of the acceptor^43^. Values from single CCSs (Fig S3H) were consistent with those from whole plasma membrane (Fig S3, A-C).

However, in general, nearly 10,000 photon counts are required to fit a bi-exponential^44, 71^. So, curve-fitting on the fluorescence decay curves from very small CCSs will not be adequate. Thus, instead to estimate FRET efficiency through curve-fitting, mean fluorescence lifetime, the center mass of the fluorescence lifetime decay, was used as an indicator for FRET efficiency (Fig 1D and Fig S3D).

For FLIM measurements with dipicrylamine (DPA; City Chemical LLC), a 20 mM stock solution of DPA in DMSO (Sigma-Aldrich, D2650) was prepared fresh from powder every day and diluted to a final concentration with PBS. Imaging was performed at least 10 min after addition of DPA. FRET efficiency between DPA and miRFP should be very low because DPA absorbance spectra and miRFP emission spectra are sufficiently separated (Fig S2C).

For live cell FLIM imaging (Fig S6), cells were imaged in imaging buffer (130 mM NaCl, 2.8 mM KCl, 5 mM CaCl_2_, 1 mM MgCl_2_, 10 mM HEPES, and 10 mM glucose at pH 7.4) at 21°C.

### Platinum replica EM

EM samples were prepared as described previously^4, 69^. Coverslips were transferred from glutaraldehyde into 0.1% w/v tannic acid for 20 min. Then, they were rinsed 4 times with water, and placed in 0.1% w/v uranyl acetate for 20 min. The coverslips were then dehydrated, critical point dried with a critical point dryer (Tousimis Samdri, 795), and coated with platinum and carbon with a Freeze fracture (Leica, EM ACE 900). The region of interest on the coverslip marked by a diamond scriber was imaged with a 20× phase-contrast objective to obtain another map of the region imaged in fluorescence. The replicas were lifted and placed onto Formvar/carbon-coated 75-mesh copper TEM grids (Ted Pella, 01802-F) that were freshly glow-discharged with PELCO easiGlow 91000. Again, the grid was imaged with a 20× phase-contrast objective to find the same region that was originally imaged in fluorescence. Each cell of interest was located on the grid prior to EM imaging^39^. TEM imaging was performed as previously described^36^ at ×15,000 magnification (1.2 nm per pixel) using a JEOL 1400 and SerialEM freeware for montaging^72^. Electron microscopy montages were processed using IMOD freeware^73^.

### FRET-CLEM image analysis

The FLIM images were aligned to the EM images using an affine spatial transformation with nearest-neighbor interpolation to map the CCSs visible in both FLIM images and EM images^35, 40^. Since position of sphere clathrin is sometimes changed during critical point drying due to a weak attachment to the membrane, we used flat and domed clathrin as fiducials^69^. Rectangular ROI was created around single isolated CCS, and mean fluorescence lifetime within each ROI was calculated. These values were analyzed by categorizing according to lattice structures, and were compared to average value of flat clathrin for each unroofed membrane. This is because FRET efficiencies, and thus fluorescence lifetimes, varied among cells due to the differences in expression levels of doner and acceptor probes, or incorporated density of DPA (Figs S4, S5 and S7).

For FRET between EGFP and ShadowY, because ShadowY expression cannot be confirmed with fluorescence, only cells with cellular average fluorescence lifetime of less than 2.1 ns were analyzed to ensure expression of ShadowY.

### EM image analysis

Binary masks of the flat (no visible curvature), domed (curved but can still see the edge of the lattice), and sphere clathrin (curved beyond a hemisphere such that the edge of the lattice is no longer visible) were manually segmented (Fig S14A)^42, 74^. The percentage of occupied membrane area was defined as the sum of areas from clathrin lattices of the specified subtype divided by the total area of measured membrane. Transient expression of CLC probes did not change the size of clathrin lattice structures but changed the density of CCS slightly (Fig S14, B-D).

### Transferrin uptake assay

HeLa cells were incubated in starvation medium [DMEM containing 20 mM HEPES at pH 7.4 and 0.1% w/v bovine serum albumin (Fisher Bioreagents, BP9703)] for 1h in an CO_2_ incubator. They were then incubated with 500 nM AP21967 or 0.1% w/v ethanol (control) in starvation medium for 5min. Then, 25 mg/mL Alexa Fluor 647 conjugated human transferrin (Invitrogen, T23366) was added to the starvation medium^75^. After 15 min incubation, the cells were fixed with 2% paraformaldehyde at room temperature for 25 min. They then washed with PBS. The cells were imaged with a Leica Falcon SP8 confocal microscope with a 63×, 1.40 NA oil immersion objective. Images were collected at a lateral spatial resolution of 120 nm per pixel. The confocal aperture was set at a diameter of 191 μm, and optical sections with z-spacing of 0.8 μm were collected. Alexa Fluor 647 was excited by 633 nm and 638-800 nm fluorescence was collected.

The stack images were recombined using a sum-intensity operation. And background intensity from non-cell regions were subtracted. Then, cell outlines were traced manually and the mean intensity (total intensity/area) was measured for individual cells. Fluorescence intensity was normalized by the average value of non-transfected cells for each coverslip.

### Live cell TIRF imaging and analysis

Images were acquired using a Nikon Eclipse Ti inverted microscope with a 100×, 1.49 NA objective and an Andor iXon Ultra 897 EM-CCD camera. HeLa cells in the phenol red free DMEM growth medium [DMEM (Gibco, 31053036) with 10% fetal bovine serum, 1:100 dilution of 100× GlutaMAX (Gibco, 35050061), 1:100 dilution of 100 mM sodium pyruvate (Gibco, 11360070)] were mounted in a chamber at 37°C with a water bath and continuous flow of humidified 5% CO_2_ to maintain the osmolality and pH of the medium. Pixel size was 110 nm. TIRF images were acquired with 100 ms exposures at 0.5 Hz for 10 min. For AP21967 treatment, images were acquired at least 10 min after addition of 500 nM AP21967.

FKBP-EGFP-CLC spots were tracked using the ImageJ plugin TrackMate^76^. Spot detection was done with the difference of Gaussians approach, and tracking was done using the simple linear assignment problem (LAP) tracker algorithm with a linking maximum distance of 330 nm per frame. Tracks that appeared and disappeared during the whole movie with over 20 s duration, and that total net displacement was less than 330 nm were selected for residence time analysis. Further, each track was visually inspected for isolation and tracking errors.

### FRET simulation

FRET simulations (Fig S1D and S13) were performed using MATLAB. We used 60 Å for Förster radius for EGFP-ShadowY FRET. For the simulation in Fig S1D, we determined the position of N- and C-terminal positions of CLC according to the structure model (PDB 3LVG and 6WCJ)^17, 45^. For the simulation in Fig S13, lateral position of N-terminus was moved by 0.6 Å spacing, and N-terminus was assumed to locate axially 25 Å higher than C-terminus. FRET efficiency between EGFP-CLC and surrounding five CLC-ShadowY, or FRET efficiency between EGFP-CLC and opposite ShadowY-CLC was calculated at different expression levels in ShadowY probes (CLC-ShadowY or ShadowY-CLC).

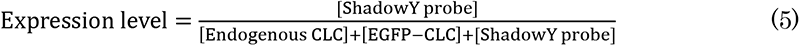

### Quantification and Statistical Analysis

Image analysis and quantification were performed with ImageJ^77^ and MATLAB. The statistical tests used for each experiment and the exact sample numbers (*n* values) are indicated in the corresponding figure legends. p values of < 0.05 were considered statistically significant. All statistical analysis and fitting were performed using Origin 2016 (Origin Lab).

## Acknowledgments

We thank Dr. L.D. Islas (Universidad Nacional Autónoma de México) and Dr. H. Murakoshi (National Institute for Physiological Sciences) for sharing the spectra of DPA and ShadowY respectively. We thank US National Heart, Lung, and Blood Institute Light Microscopy Core and Electron Microscopy Core for use of instruments and advice. We thank G. Haber for help with coding. We thank member of the Taraska lab for helpful discussions and edits. J.W.T. is supported by the Intramural Research Program of the National Heart Lung and Blood Institute, National Institutes of Health. K.O. is supported by JSPS Research Fellowship for Japanese Biomedical and Behavioral Researchers at NIH.

## Author contributions

K.O., K.A.S., and J.W.T. designed the research. K.O. performed experiments and analysis. K.A.S. developed software for analysis. M-P.S. helped molecular cloning. J.W.T. supervised the project. K.O. and J.W.T. wrote the paper. All authors contributed to the interpretation of the data and commented on the manuscript.

## Supplementary figure legends

**Figure S1.**
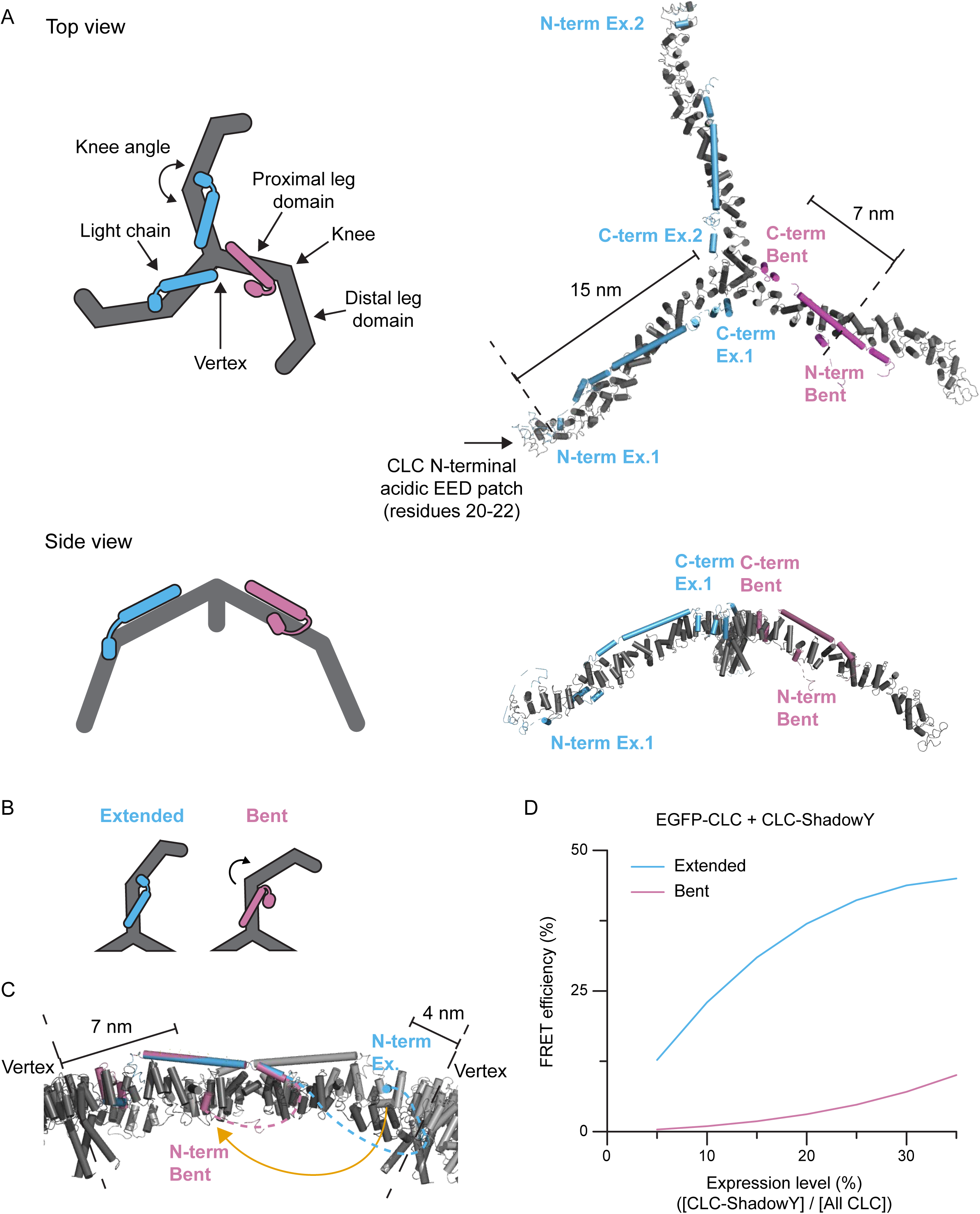
Structural model of clathrin triskelion. (A) A structural model of a clathrin triskelion. Top and side views are shown. The model is based on PDB 3LVG. (B) Schematic models for conformational changes in CLC and heavy chain knee. (C) Side view of Fig 2B. The distances between triskelion vertex (CLC C-terminus) and N-terminus of extended or bent CLC are shown. (D) Simulations on FRET efficiency between EGFP-CLC and near five CLC-ShadowY at different CLC-ShadowY expression levels.

**Figure S2.**
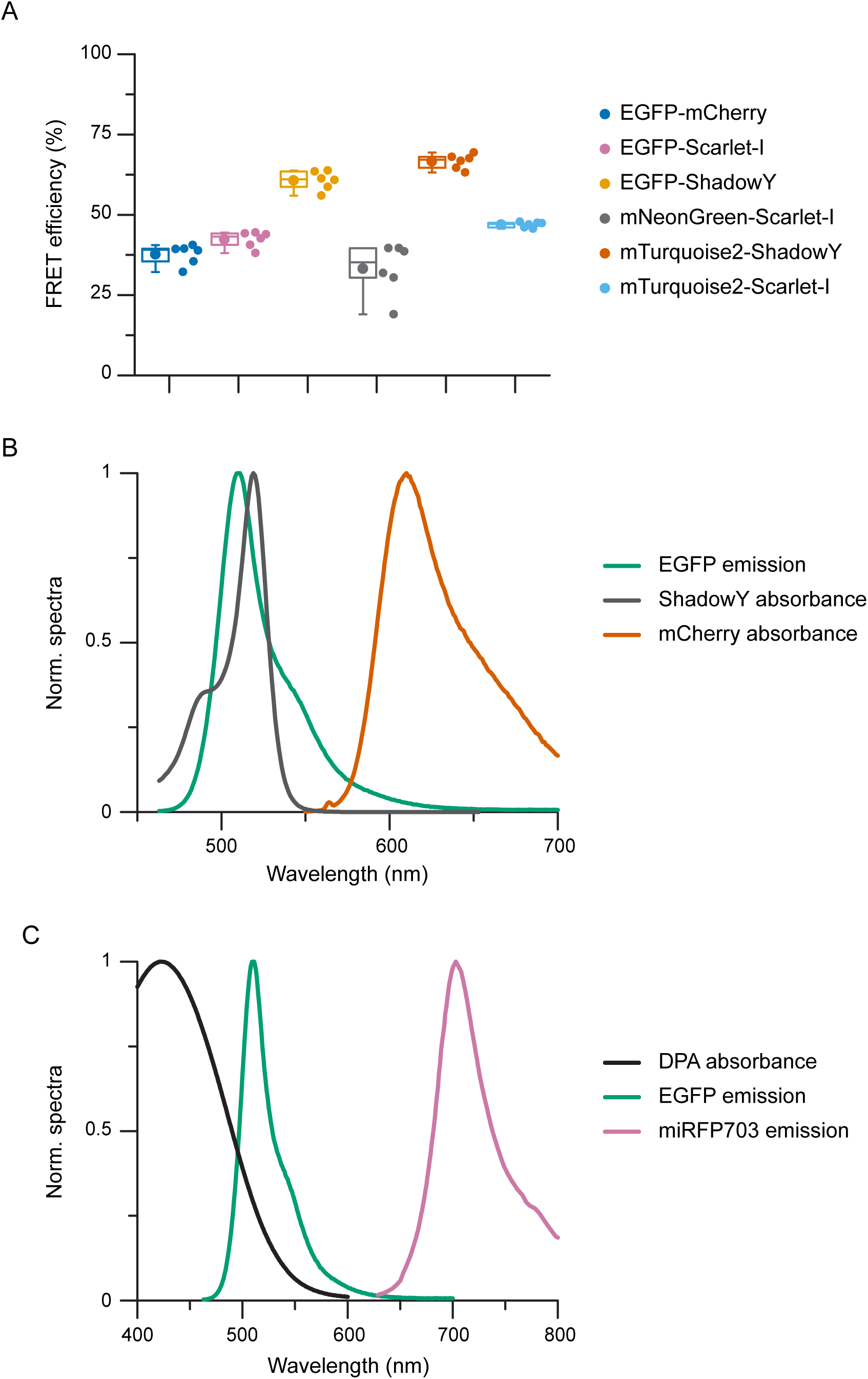
FRET pairs used in this study. (A) FRET efficiencies of tandemly-connected fluorescent proteins (FPs). Fluorescence lifetimes were measured in fixed HeLa cells expressing FPs or tandemly-connected FPs. Amplitude-weighted fluorescence lifetimes were estimated from fluorescence lifetime decays by fitting with a bi-exponential. FRET efficiencies were then calculated from amplitude-weighted fluorescence lifetimes of the donor only and tandem-connected FPs. *n* = 6 cells for each condition. (B) Normalized spectra of emission of EGFP, absorbance of ShadowY, and absorbance of mCherry. (C) Normalized spectra of absorbance of DPA, emission of EGFP, and emission of miRFP703.

**Figure S3.**
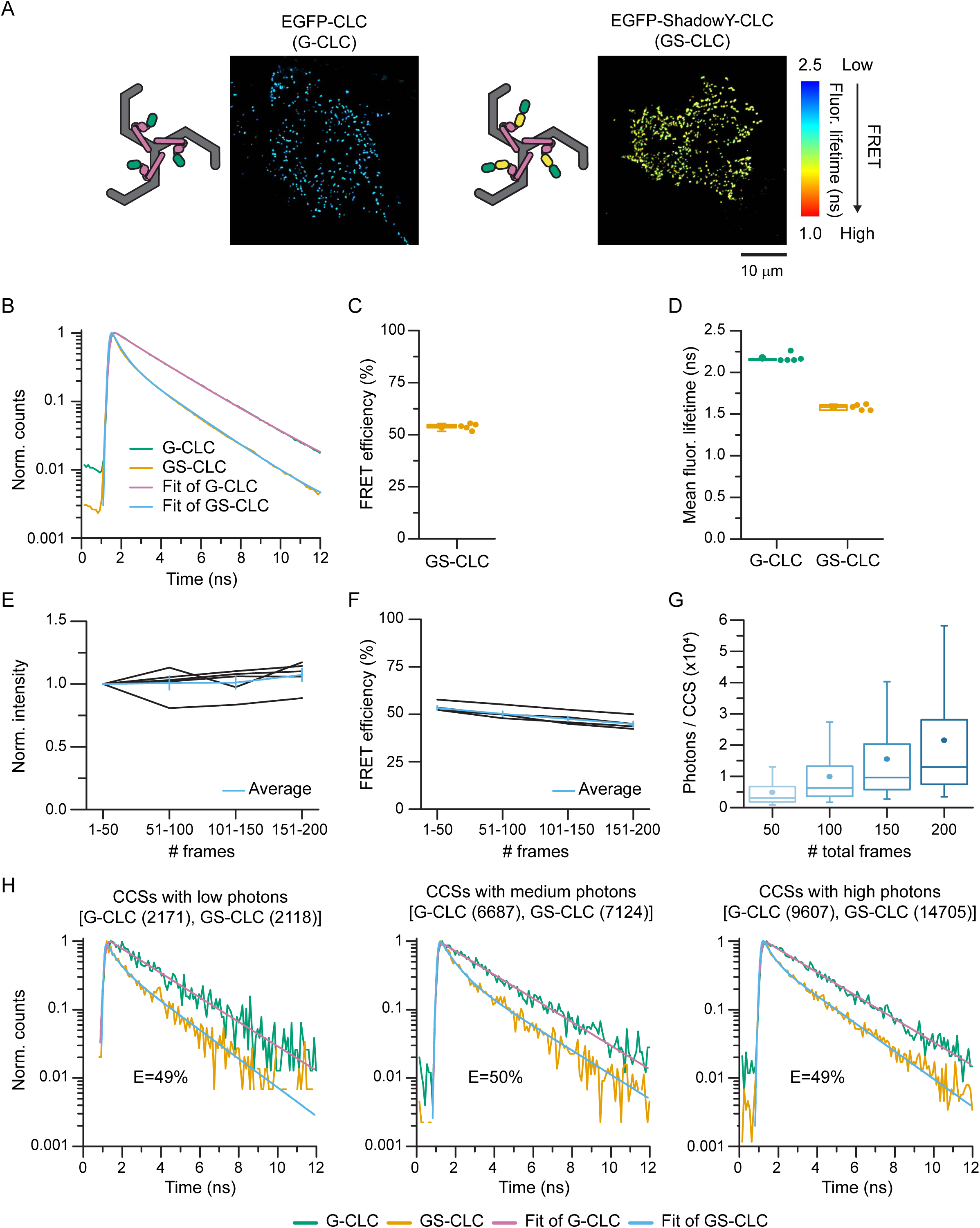
FLIM-FRET imaging with EGFP and ShadowY at single CCS resolution. (A) FLIM images of unroofed membranes of HeLa cells expressing EGFP-CLC (left) or EGFP-ShadowY-CLC (right). Scale 10 μm. (B) Fluorescence lifetime decays from EGFP-CLC or EGFP-ShadowY-CLC on unroofed membranes. And fluorescence lifetime decays were fitted with a bi-exponential. (C) FRET efficiencies of EGFP-ShadowY-CLC on unroofed membranes. *n* = 5 cells. (D) Mean fluorescence lifetimes of EGFP-CLC or EGFP-ShadowY-CLC on unroofed membranes. *n* = 5 cells for each condition. (E and F) Unroofed membranes of HeLa cells expressing EGFP-ShadowY-CLC were imaged repeatedly (50 frames, 4 times). Changes in fluorescence intensity (F) and FRET efficiency (G). *n* = 5 cells. (G) Histogram of photon counts per single CCSs on an unroofed membrane of a HeLa cell expressing EGFP-ShadowY-CLC with medium level expression. *n* = 322 CCSs from 1 cell. (H) Representable fluorescence lifetime decays of single CCSs on unroofed membranes of HeLa cells expressing EGFP-CLC or EGFP-ShadowY-CLC with different total photon counts indicated in brackets. The fluorescence lifetime decays were fitted with a bi-exponential and FRET efficiencies were estimated.

**Figure S4.**
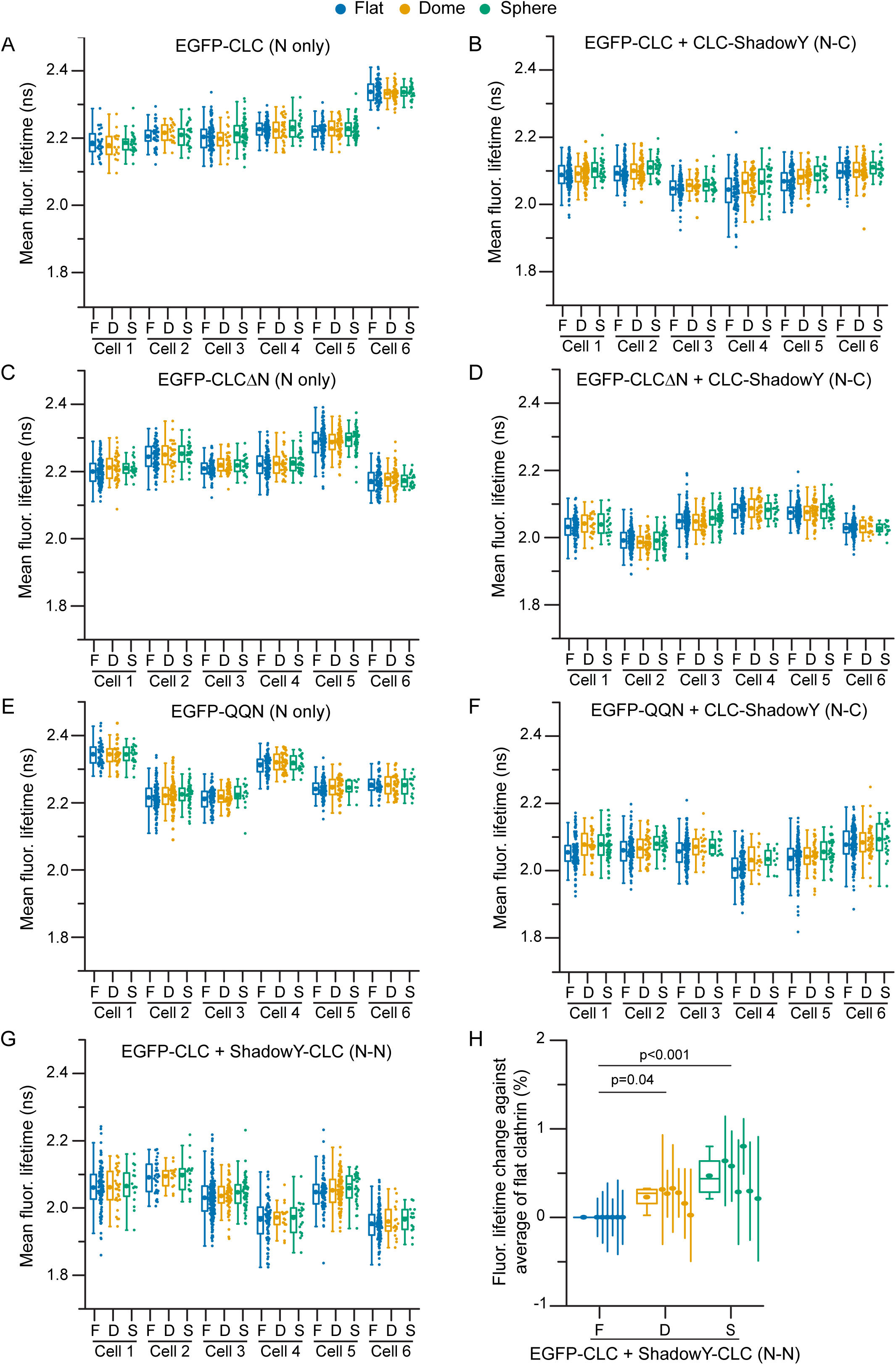
Mean fluorescence lifetimes of FRET-CLEM measurements on EGFP and ShadowY attached CLC probes, and FRET-CLEM measurements between N- and N- terminus. (A-F) Mean fluorescence lifetimes from single CCSs of experiments in Fig 2. *n* (flat, domed, sphere CCSs/cell) = (30-107, 16-49, 22-53) for EGFP-CLC (A), (82-163, 32-77, 22-32) for EGFP-CLC and CLC-ShadowY (B), (78-128, 28-76, 22-55) for EGFP-CLCΔN (C), (94-166, 18-74, 13-60) for EGFP-CLCΔN and CLC-ShadowY (D), (42-129, 26-69, 7-49) for EGFP- QQN (E), and (107-183, 15-52, 9-68) for EGFP-QQN and CLC-ShadowY (F). (G, H) FRET-CLEM was performed on HeLa cells expressing EGFP-CLC and ShadowY- CLC. Mean fluorescence lifetimes from single CCSs (G) and their changes (H) were analyzed by categorizing them according to lattice structures (flat, domed and sphere). For plot H, each dot is from one cell experiment and errors are SE. *n* = 6 cells, *n* (flat, domed, sphere CCSs/cell) = (32-202, 15-88, 15-56). One-way ANOVA, then Tukey’s test. For box plots, box is interquartile range, center line is median, center circle is mean, outliers are a coefficient value of 1.5.

**Figure S5.**
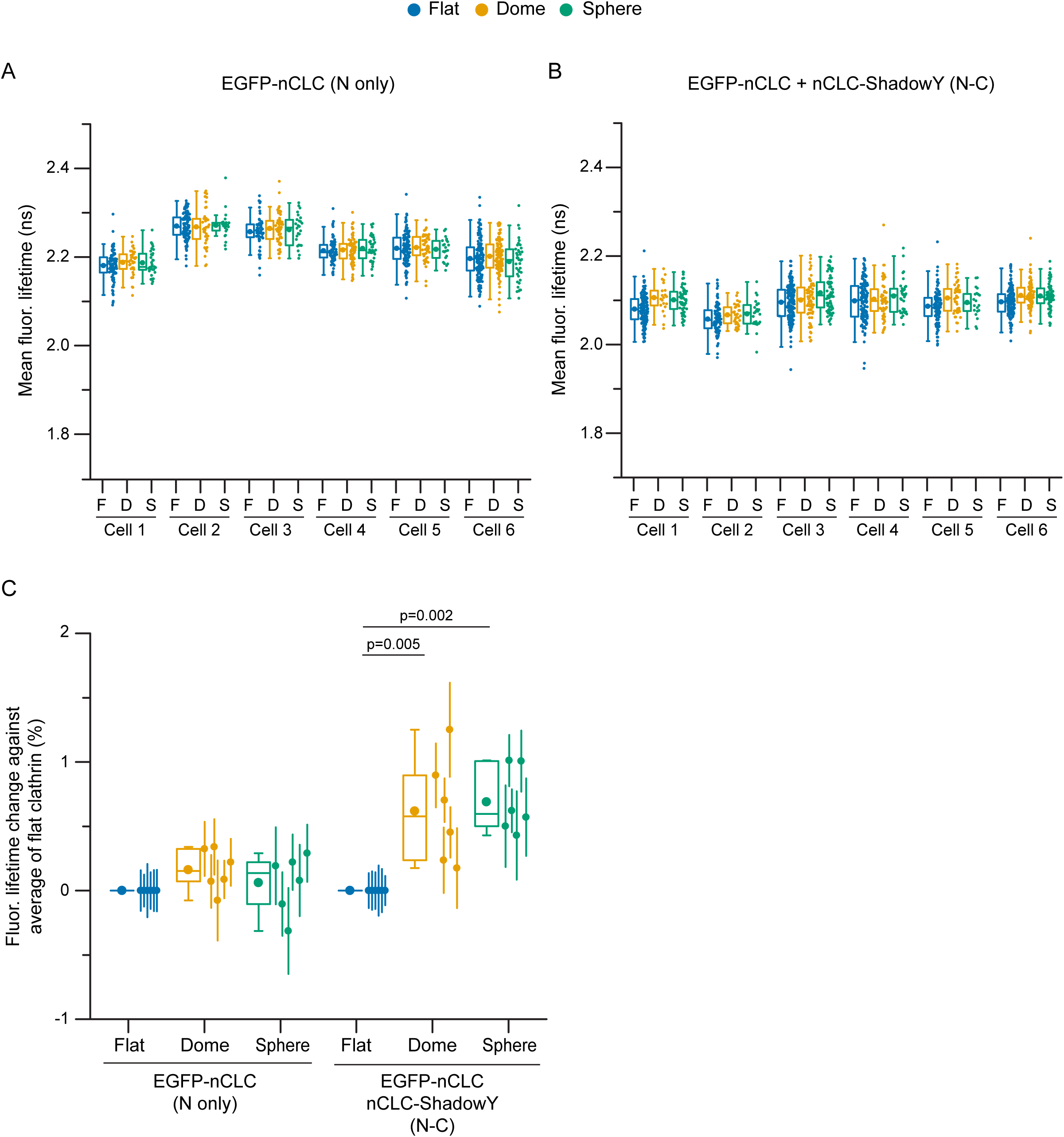
FRET-CLEM with neuronal CLC. FRET-CLEM was performed on HeLa cells expressing either EGFP-nCLC (neuronal isoform), or EGFP-nCLC and nCLC-ShadowY. Mean fluorescence lifetimes from single CCSs (A, B) and their changes (C) were analyzed by categorizing them according to lattice structures (flat, domed and sphere). For plot C, each dot is from one cell experiment and errors are SE. *n* = 6 cells each condition, *n* (flat, domed, sphere CCSs/cell) = (56-152, 38-99, 22-39) for EGFP-nCLC (A) and (105-181, 21-81, 20-71) for EGFP-nCLC and nCLC-ShadowY (B). One-way ANOVA, then Tukey’s test. For box plots, box is interquartile range, center line is median, center circle is mean, outliers are a coefficient value of 1.5.

**Figure S6.**
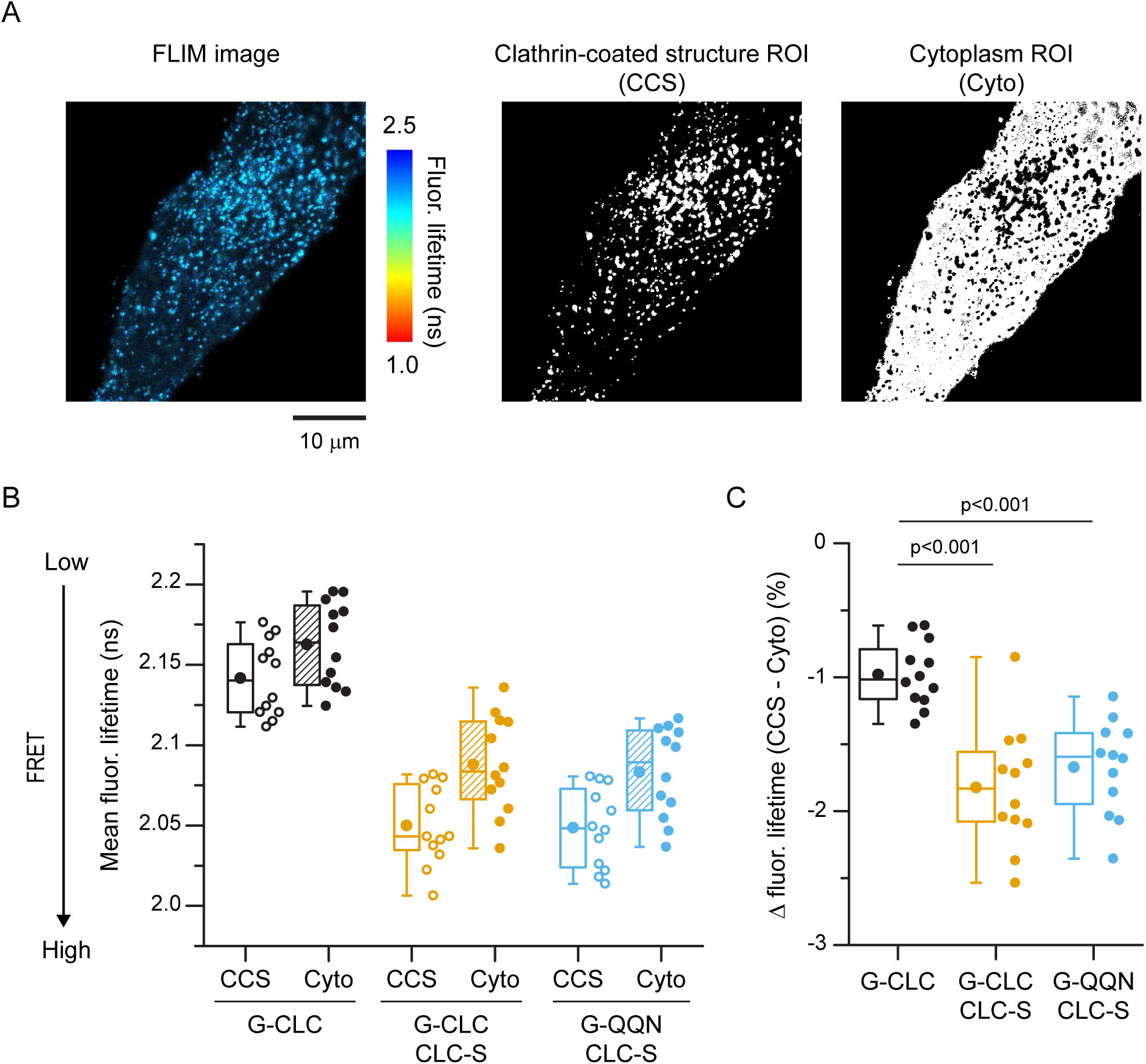
FLIM imaging in living cells. (A) A FLIM image of a living HeLa cell expressing EGFP-CLC (left). And ROIs for CCSs (CCS, center) and cytoplasm (Cyto, right) are shown. Scale 10 μm. (B) Mean fluorescence lifetimes within CCS ROI or Cyto ROI of EGFP-CLC (G-CLC), EGFP-CLC and CLC-ShadowY (G-CLC + CLC-S), or EGFP-QQN and CLC- ShadowY (G-QQN + CLC-S) expressing cells. *n* = 12 cells from 2 experiments for each conditions. (C) Differences in mean fluorescence lifetimes between CCS ROI and Cyto ROI. One-way ANOVA, then Tukey’s test. For box plots, box is interquartile range, center line is median, center circle is mean, outliers are a coefficient value of 1.5.

**Figure S7.**
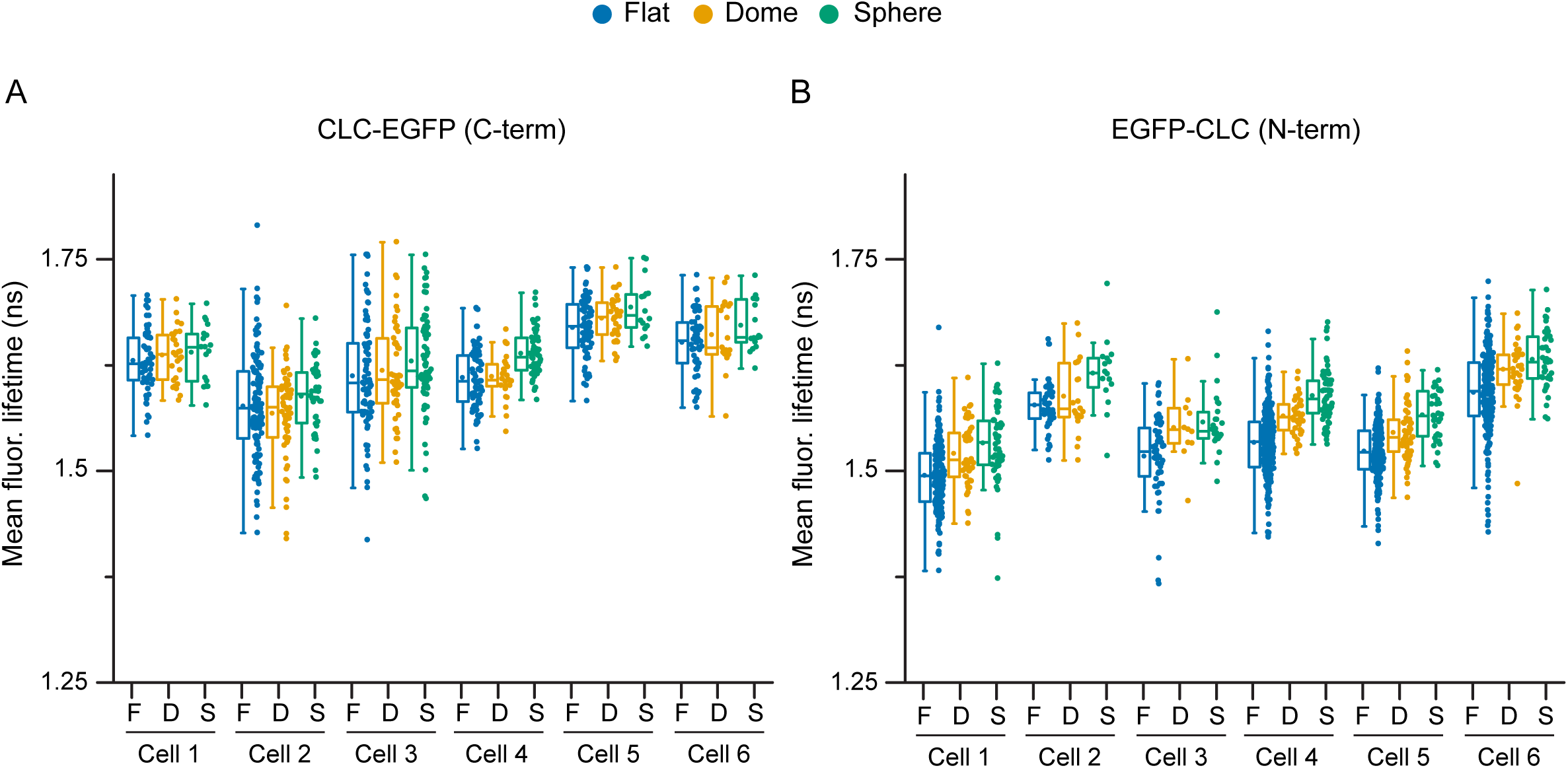
Mean fluorescence lifetimes of FRET-CLEM measurements with DPA. Mean fluorescence lifetimes from single CCSs of experiments in Figure 3E. *n* (flat, domed, sphere CCSs/cell) = (45-239, 11-59, 16-65) for CLC-EGFP (A) and (52-128, 19-58, 16-72) for EGFP-CLC (B).

**Figure S8.**
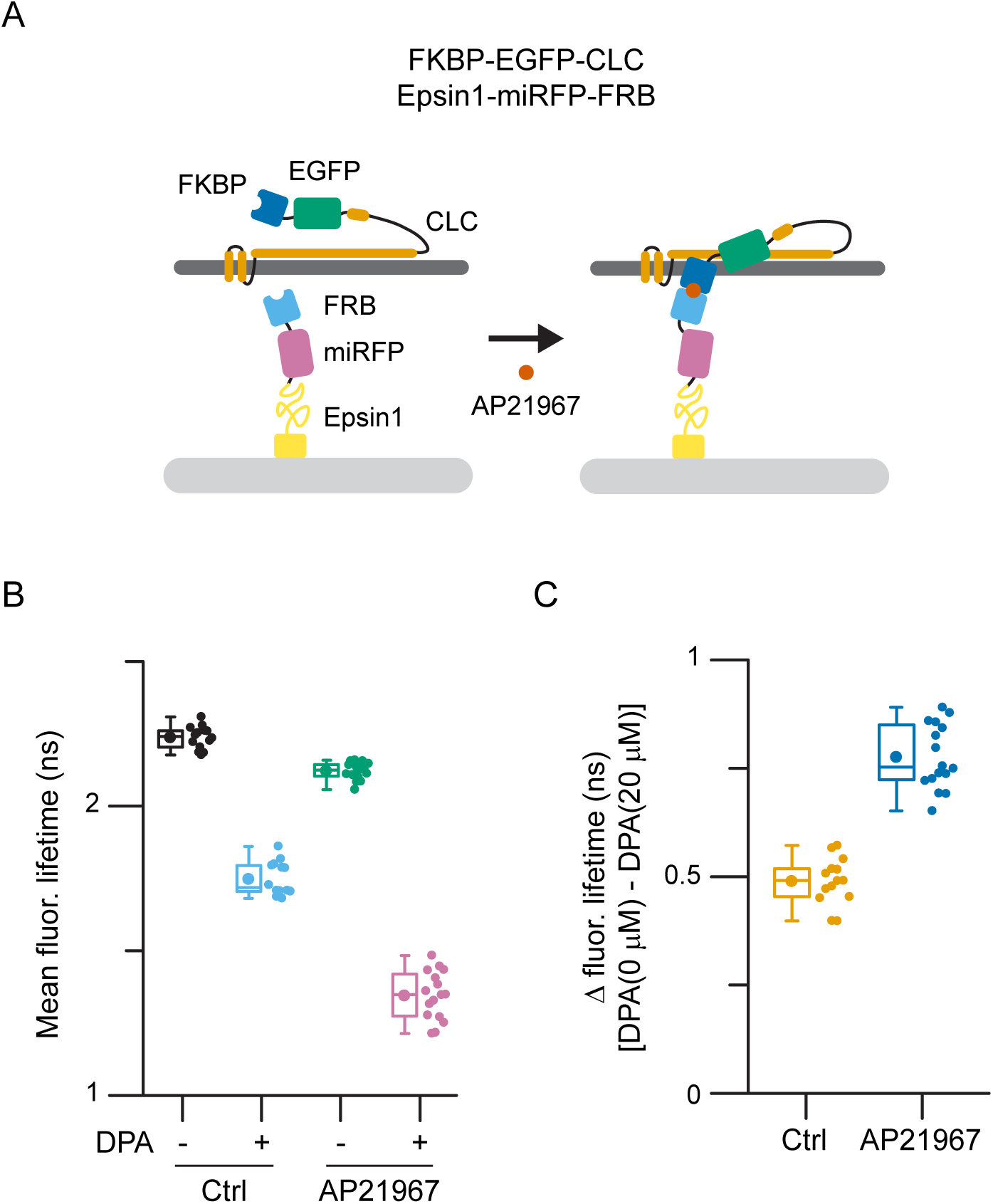
Manipulation of CLC N-terminal position with epsin1-FRB construct. (A) Schematic models of the chemically inducible FKBP/FRB dimerization system. FKBP is attached to the N-terminus of CLC, and FRB is attached to the C-terminus of epsin1. A rapamycin analog, AP21967, induces heterodimerization between FKBP and the T2098L mutant of FRB. (B) Fluorescence lifetime measurements were performed on unroofed membranes of HeLa cells expressing FKBP-EGFP-CLC and epsin1-miRFP-FRB without or with 20 μM DPA. Cells were unroofed after 15 min incubation with AP21967 or ethanol (control). *n* = 14 (control), and 16 cells (AP21967) from 2 experiments. (C) Differences in fluorescence lifetimes without and with DPA.

**Figure S9.**
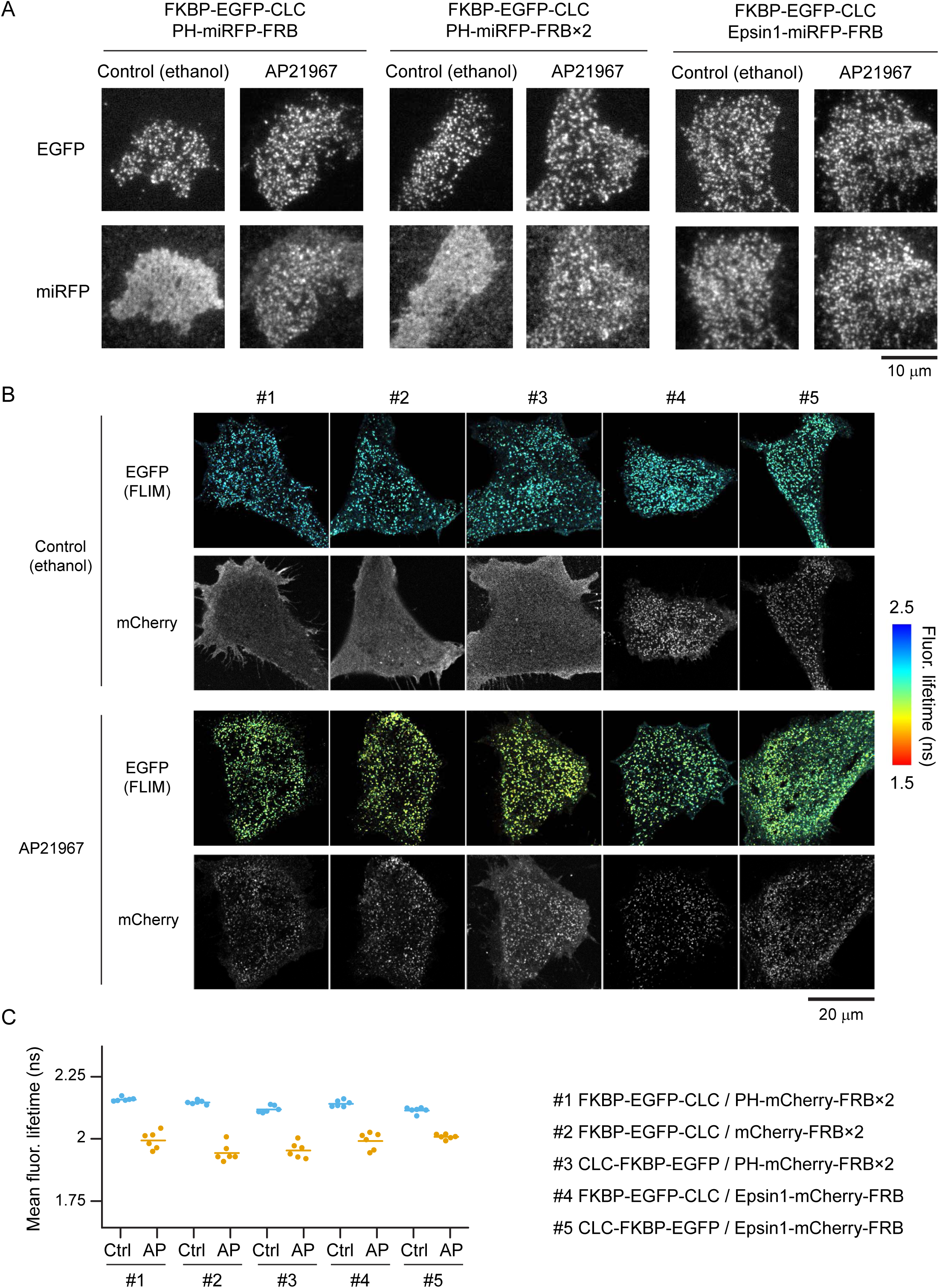
FKBP/FRB dimerization was confirmed by changes in probe distributions and FRET efficiencies. (A) TIRF images of unroofed membranes of HeLa cells expressing FKBP-EGFP-CLC and either PH-miRFP-FRB (left), PH-miRFP-FRB×2 (center), or Epsin1-miRFP- FRB (right). Cells were unroofed after 15 min incubation with AP21967 or ethanol (control). Scale 10 μm. (B) FLIM and confocal images of fixed HeLa cells expressing FKBP and FRB probes treated with AP21967 or ethanol (control). Scale 20 μm. (C) Mean fluorescence lifetimes of fixed HeLa cells expressing FKBP and FRB probes treated with AP21967 or ethanol (control). *n* = 6 cells for each conditions.

**Figure S10.**
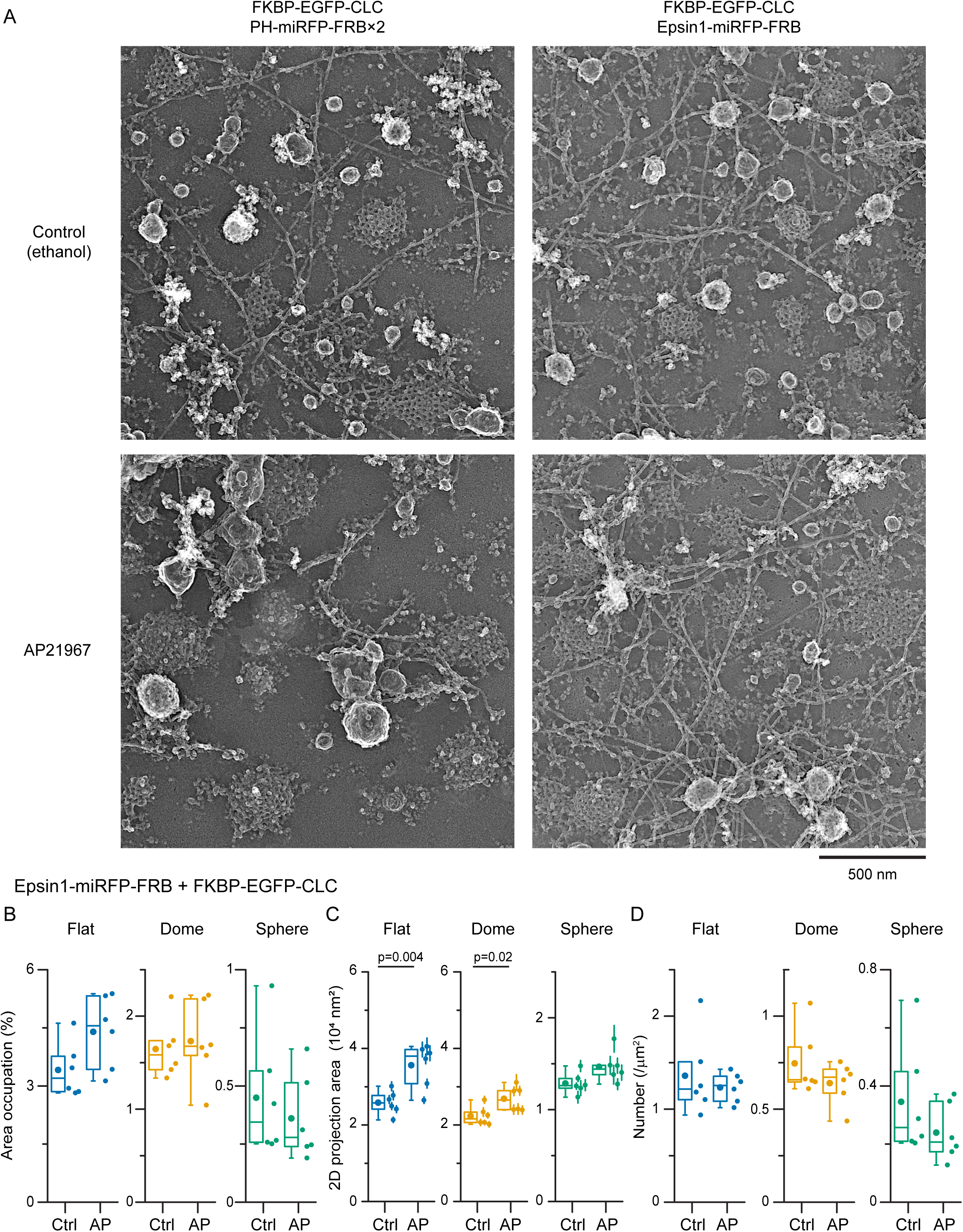
PREM analysis on lattice structures with manipulation of CLC conformation. (A) PREM images of unroofed membranes of HeLa cells expressing FKBP-EGFP-CLC with either PH-miRFP-FRB×2 (left) or Epsin1-miRFP-FRB (right). Cells were treated with ethanol (control, top) or AP21967 (bottom) for 15 minutes before unroofing. Scale 500 nm. (B-D) Unroofed membranes from cells expressing FKBP-EGFP-CLC and epsin1- miRFP-FRB treated with AP21967 or ethanol (control) were imaged with PREM. Two-dimension area of single CCS were manually segmented and measured. Membrane area occupation against the total analyzed membrane area (B), two-dimension projection area (C), and density (D) of flat, domed, and sphere CCSs were compared. Each dot is from one cell experiment. *n* = 6 cells for each condition. The average measured area (mean ± SE) = 144 ± 19 (control) and 139 ± 13 μm^2^ (AP21967). For box plots, box is interquartile range, center line is median, center circle is mean, outliers are a coefficient value of 1.5. An unpaired t test was used.

**Figure S11.**
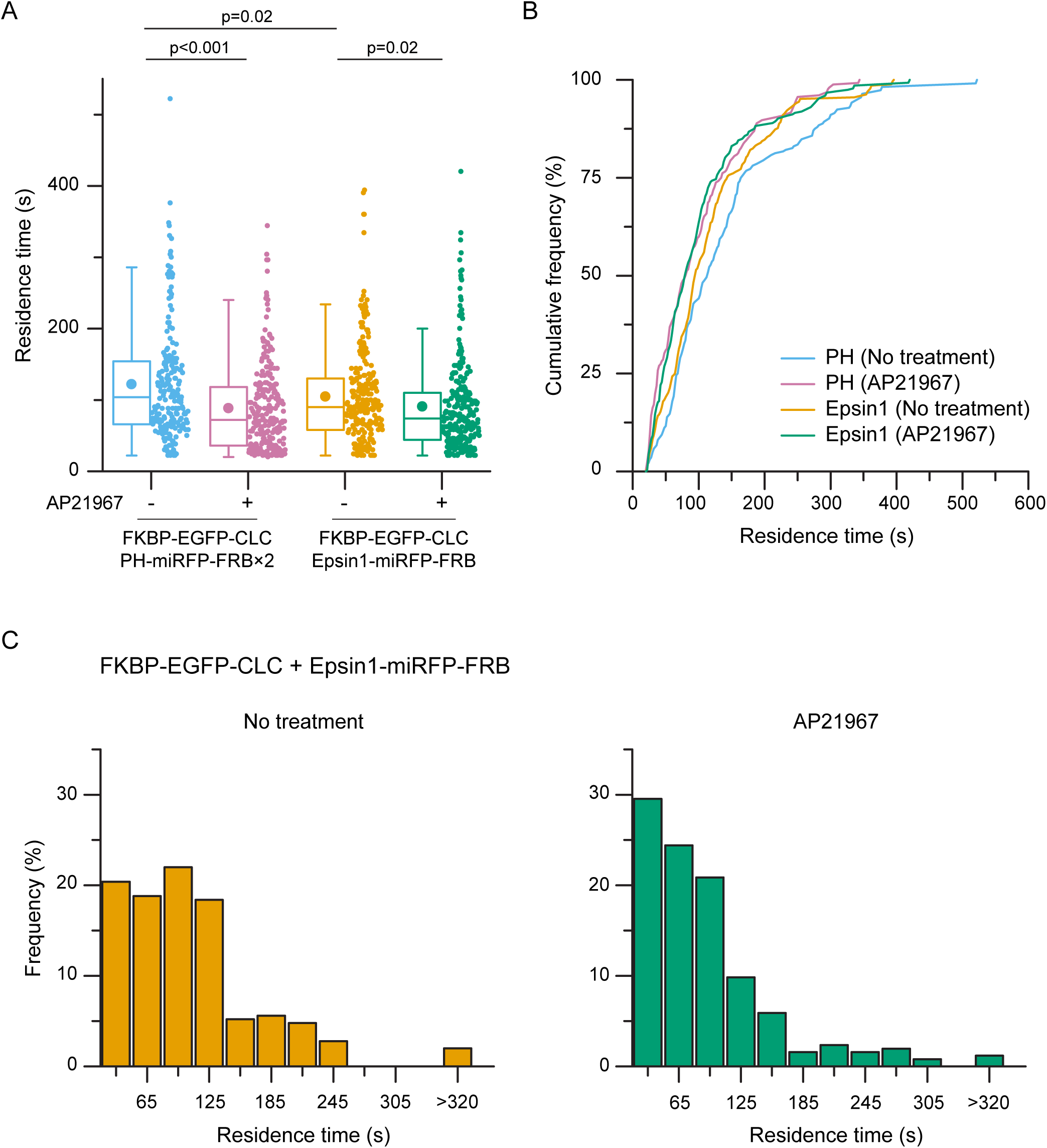
Live cell TIRF imaging of CCSs with manipulation of CLC conformation. Lice cell TIRF imaging was performed with HeLa cells expressing FKBP-EGFP-CLC and either PH-miRFP-FRB×2 (shown in Fig 5, D and E) or epsin1-miRFP-FRB without or with AP21967. Tracks with over 20 s residence time were analyzed. (A) Residence times of tracked spots. *n* = 250 spots from 5 cells (no treatment) and 255 spots from 5 cells (AP21967) for epsin1-miRFP-FRB. For box plots, box is interquartile range, center line is median, center circle is mean, outliers are a coefficient value of 1.5. An unpaired t test was used. (B) Cumulative distributions of residence times. (C) Histogram of residence times for epsin1-miRFP-FRB experiments.

**Figure S12.**
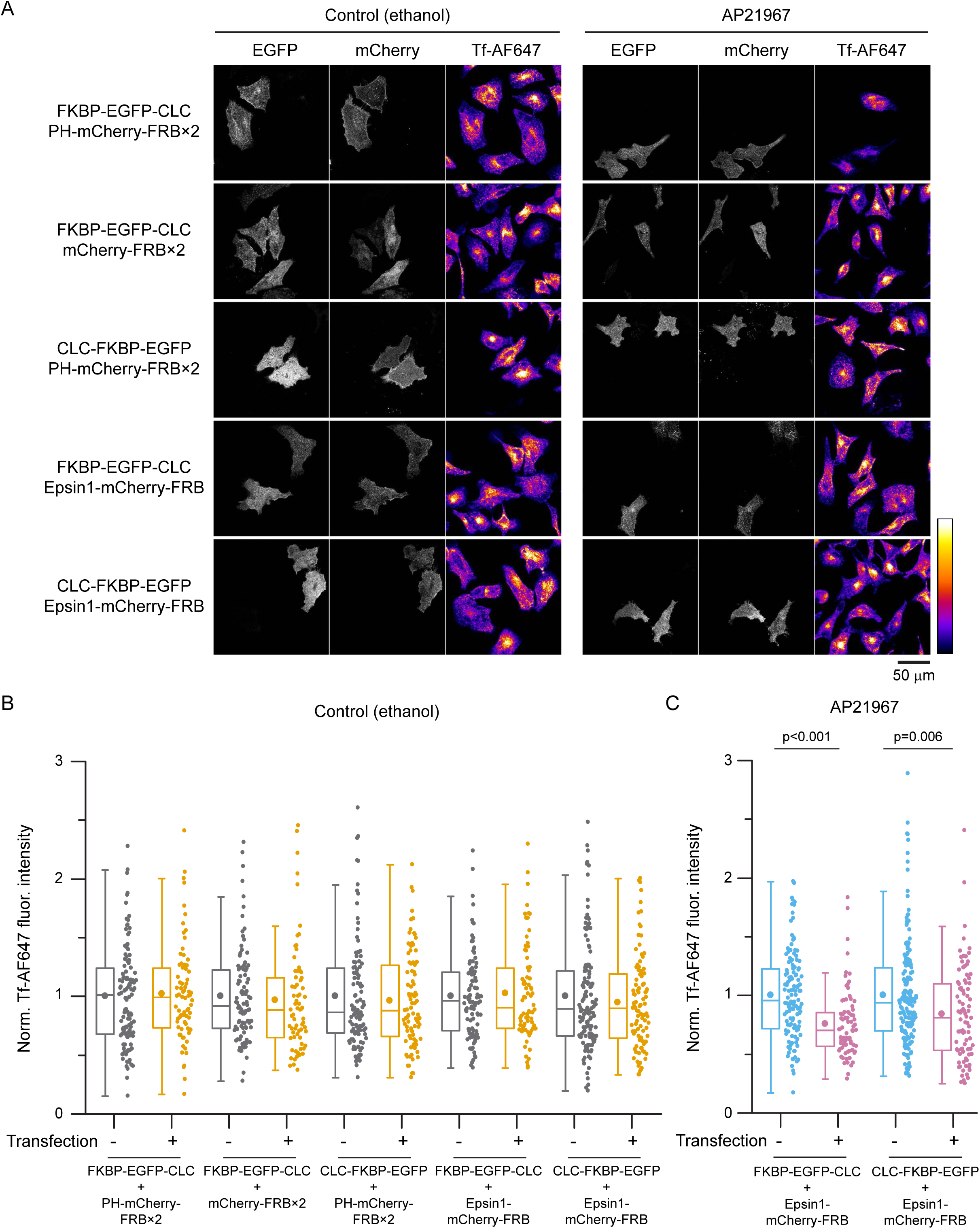
Transferrin uptake assay. (A) Confocal projection images of EGFP, mCherry, and Alexa Fluor 647 conjugated transferrin (Tf-AF647) in HeLa cells expressing FKBP and FRB probes treated with AP21967 or ethanol (control). Fluorescence intensity of Tf-AF647 is represented by pseudo color. Scale 50 μm. (B) Transferrin uptake in HeLa cells expressing FKBP and FRB probes treated with ethanol (control). Fluorescence intensities of incorporated Tf-AF647 normalized by non-transfected cells in the same sample were compared between non-transfected and transfected cells. *n* = 70-129 cells from 2 experiments for each conditions. (C) Transferrin uptake in HeLa cells expressing epsin1-mCherry-FRB with FKBP-EGFP-CLC or CLC-FKBP-EGFP treated with AP21967. *n* = 78-173 cells from 2 experiments for each conditions. For box plots, box is interquartile range, center line is median, center circle is mean, outliers are a coefficient value of 1.5. An unpaired t test was used.

**Figure S13.**
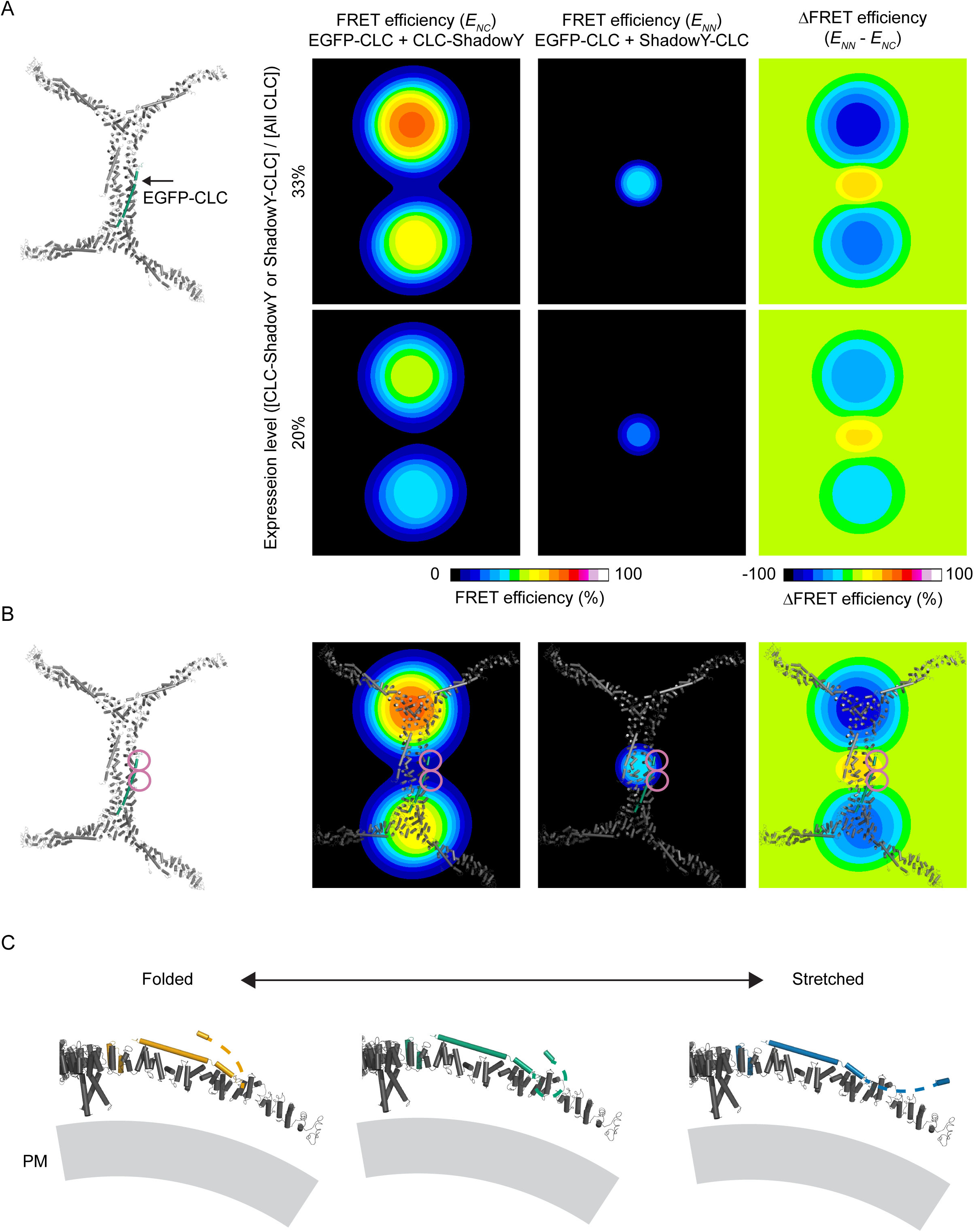
FRET simulations at various CLC N-terminal positions in relation the heavy chain proximal leg domain. (A) Simulations of FRET efficiency between EGFP-CLC (indicated by arrow in structural model) and near five CLC-ShadowY (left column; *E_NC_*), or between EGFP-CLC and opposite ShadowY-CLC (center column; *E_NN_*). FRET efficiencies were calculated by moving N-terminal position of EGFP-CLC at 33% (top) or 20% (bottom) expression levels in ShadowY probes. Right column shows the differences in FRET efficiencies between *E_NN_* and *E_NC_*. The model is based on PDB 3LVG and 6WCJ. 3LVG is overlaid with 6WCJ. (B) Structural models are overlaid with simulation results at 33% expression level. Magenta circles indicate the expected N-terminal position of CLC. (C) Structural models of possible CLC conformation with various folding states in N-terminal domain. The model is based on PDB 3LVG.

**Figure S14.**
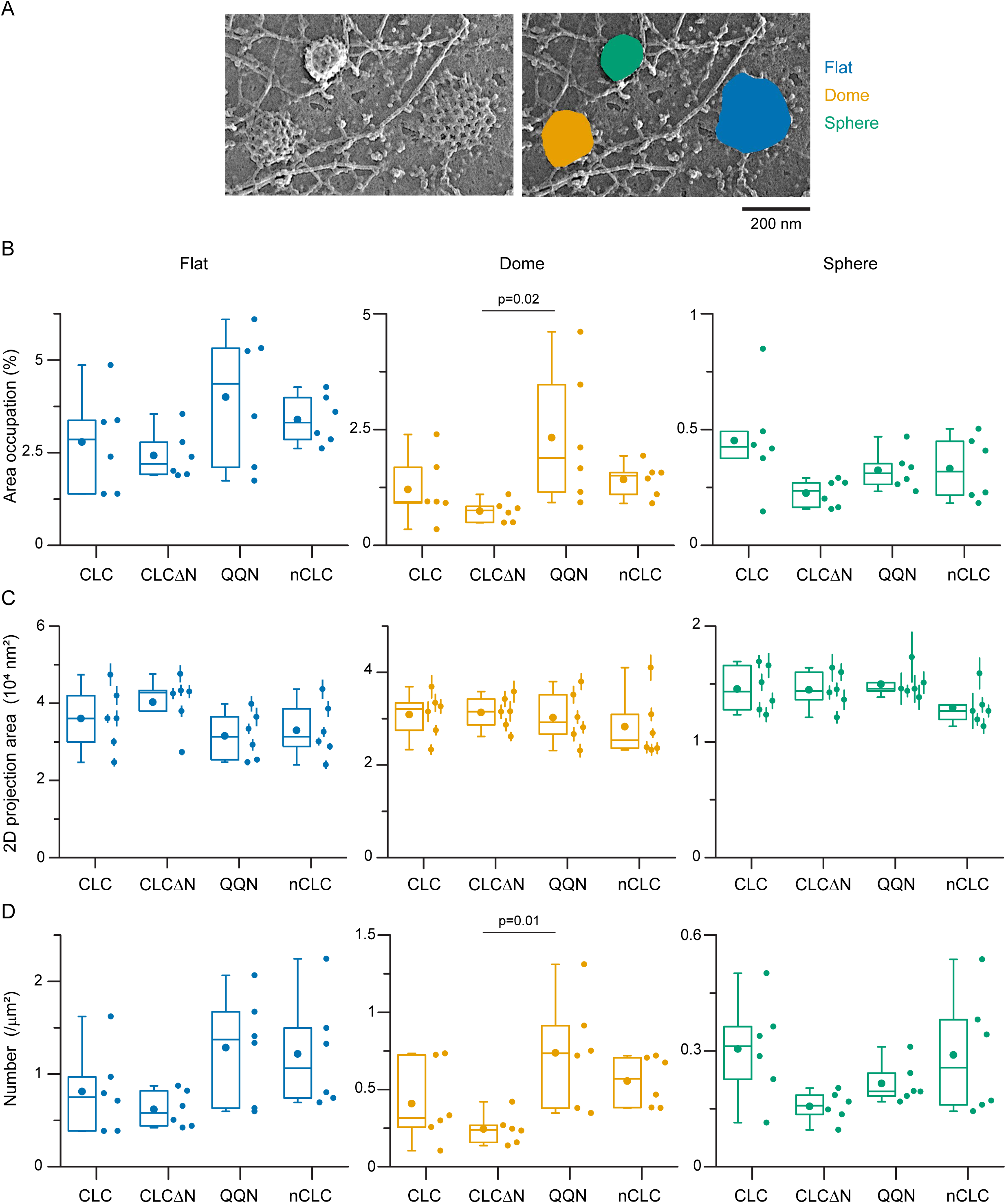
PREM analysis on lattice structures. PREM data from Fig 2, D to F and Fig S5 were analyzed. Two-dimension area of single CCS were manually segmented and measured. *n* = 6 cells for each condition. The average measured area (mean ± SE) = 206 ± 43 (EGFP-CLC), 312 ± 26 (EGFP-CLCΔN), 143 ± 32 (EGFP-QQN), and 159 ± 27 μm^2^ (EGFP-nCLC). (A) A PREM image of an EGFP-CLC expressing cell. Right image shows segmentation example. Scale 200 nm. (B-D) Membrane area occupation against the total analyzed membrane area (B), two-dimension projection area (C), and density (D) of flat, domed, and sphere CCSs were compared. One-way ANOVA, then Tukey’s test. Each dot is from one cell experiment and errors are SE. For box plots, box is interquartile range, center line is median, center circle is mean, outliers are a coefficient value of 1.5.

---

**Table S1.**
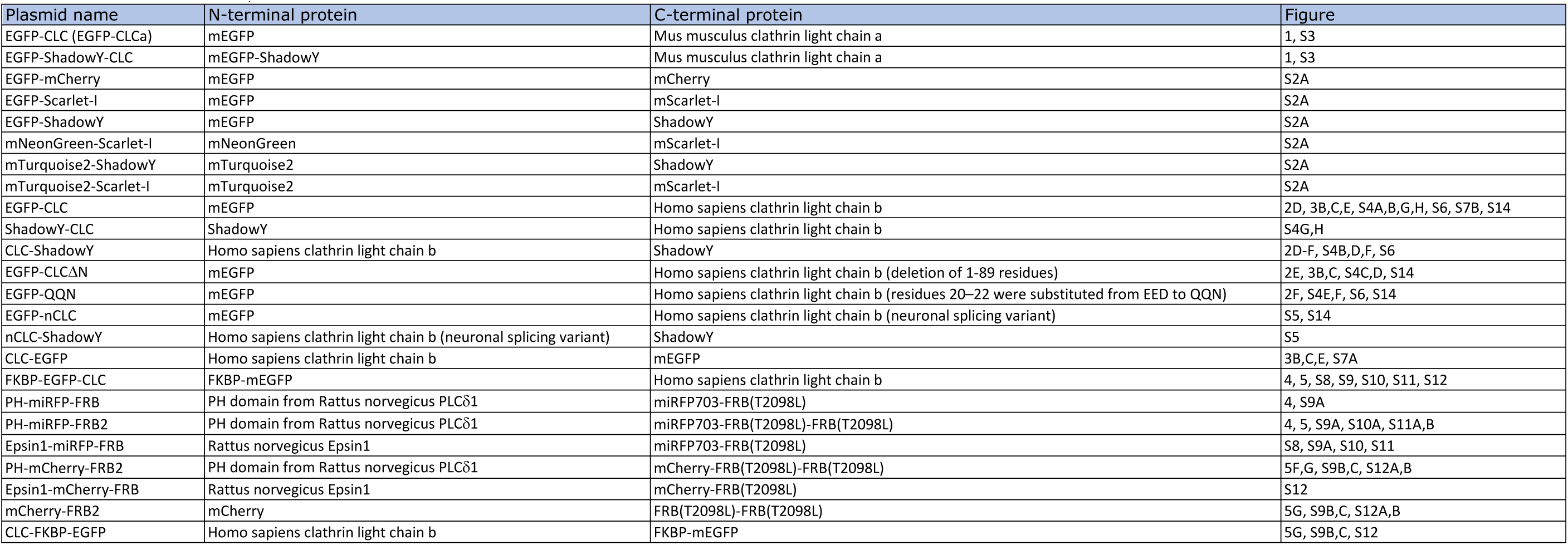
_Plasmid list used in this study._

